# Multi-omics analysis identifies drivers of protein phosphorylation

**DOI:** 10.1101/2022.06.03.494740

**Authors:** Tian Zhang, Gregory R. Keele, Isabela Gerdes Gyuricza, Matthew Vincent, Catherine Brunton, Timothy A. Bell, Pablo Hock, Ginger D. Shaw, Steven C. Munger, Fernando Pardo-Manuel de Villena, Martin T. Ferris, Joao A. Paulo, Steven P. Gygi, Gary A. Churchill

## Abstract

Phosphorylation of proteins is a key step in the regulation of many cellular processes including activation of enzymes and signaling cascades. The abundance of a phosphorylated peptide (phosphopeptide) is determined by the abundance of its parent protein and the proportion of target sites that are phosphorylated. We quantified phosphopeptides, proteins, and transcripts in heart, liver, and kidney tissue samples of mice from 58 strains of the Collaborative Cross strain panel. We mapped ∼700 phosphorylation quantitative trait loci (phQTL) across the three tissues and applied genetic mediation analysis to identify causal drivers of phosphorylation. We identified kinases, phosphatases, cytokines, and other factors, including both known and potentially novel interactions between target proteins and genes that regulate site-specific phosphorylation. Our analysis highlights multiple targets of pyruvate dehydrogenase kinase 1 (PDK1), a regulator of mitochondrial function that shows reduced activity in the NZO/HILtJ mouse, a polygenic model of obesity and type 2 diabetes.

## Introduction

Protein phosphorylation is a reversible post-translational modification (PTM) and one of the most common mechanisms for regulating protein activity and transmitting signals in cell biology [1-3]. Phosphorylation occurs at specific sites within a protein where kinases and phosphatases add and remove phosphate moieties [4]. The level of activity of kinases and phosphatases is determined by their abundance [5, 6], intracellular and extracellular stimuli [7-10], interaction with co-factors [11], and PTMs including phosphorylation [12-14]. Therefore, the phosphorylation level of a given site within a protein depends on multiple factors, any of which could be influenced by genetic variation [15, 16].

Genetic variants that affect quantitative phenotypes can be identified through quantitative trait locus (QTL) mapping in humans and in model organisms. In addition to clinical phenotypes, QTL mapping can be applied to molecular traits such as gene expression [17-22], chromatin accessibility [23], and protein abundance [24, 25]. QTL mapping of transcripts (eQTL) and proteins (pQTL) has revealed how genetic variants can alter the regulatory flow from encoded gene through transcription and translation [20, 26, 27]. However, only limited research has been conducted on how genetic variation influences protein phosphorylation or other PTMs [18, 28, 29].

Genetically diverse model organism populations increase the scope and power of QTL mapping. The Collaborative Cross (CC) [30, 31] is a panel of recombinant inbred mouse strains descended from eight founder inbred strains: A/J (AJ), C57BL/6J (B6), 129S1/SvImJ (129), NOD/ShiLtJ (NOD), NZO/HlLtJ (NZO), CAST/EiJ (CAST), PWK/PhJ (PWK) and WSB/EiJ (WSB). The founder strains represent traditional laboratory as well as wild-derived strains, encompassing three subspecies of the house mouse [32, 33] and harbor ∼50 million known genetic variants [34]. The current CC panel consists of more than 60 strains that are homozygous at most loci (> 99%). The ability to use replicate animals of CC strains is an important feature of CC studies that improves QTL mapping power [23], and enables studies of response to interventions [35-37] and other applications [38].

We previously reported on the genetic regulation of protein abundance in liver of CC strains [25]. Here we expand on our earlier investigation to examine how genetic variation regulates protein phosphorylation. We used mass spectrometry analysis to quantify the proteome and phosphoproteome across three tissues (heart, kidney, and liver) from 116 mice representing female/male pairs from 58 CC strains. We performed QTL mapping to obtain pQTL and phosphorylation QTL (phQTL). In addition, we mapped the residuals of phosphopeptide abundance after regression on the abundance of the protein they derived from, *i*.*e*., the parent protein abundance, to obtain adjusted phosphopeptide QTL (adj-phQTL). This approach allowed us to differentiate between the contributions of two distinct mechanisms that determine the abundance of phosphopeptides, the abundance of its parent protein and the proportion of target sites that are phosphorylated, with the latter likely reflecting the activity of a catalyst intermediate. We then applied mediation analysis and identified candidate genes that influence phosphorylation levels through the second mechanism.

## Results

### Quantitative phosphoproteome profiling of heart, kidney, and liver in CC mice

Heart, kidney, and liver tissue samples were collected from 116 mice representing one male and one female from each of 58 CC strains (**Table S1**). We utilized a tandem mass tag (TMT)-based proteomics workflow (**Fig. 1A**) to quantify total protein abundance and the abundance of phosphorylated peptides (phosphopeptides). We quantified 6172, 7286, and 6558 proteins, and 4975, 4236, and 4246 non-polymorphic phosphopeptides in heart, kidney, and liver tissue, respectively. The number of proteins reported for liver differs slightly from our previous study, where we report 6798 proteins, due to differences in the preprocessing and filtering steps. Nearly 5,000 proteins were quantified in all three tissues and ∼6,500 proteins were quantified in at least two tissues (**Fig. 1B**). Fewer phosphopeptides were quantified across multiple tissues; ∼1,500 were observed in all three tissues, but the majority of phosphopeptides were observed in only one tissue (**Fig. 1C)**. The number of phosphorylation sites identified for a given protein ranged from 1 to 148 (TTN in heart), with fewer than 10 sites detected for most proteins (median = 1) (**Fig. S1A**). The abundance of most phosphopeptides was slightly correlated with the abundance of their parent proteins (median correlation: heart = 0.32, kidney = 0.36, liver = 0.40) (**Fig. S1B**). To obtain an estimate of phosphorylation that is independent of the parent protein abundance, we computed the residual of phosphopeptide abundance after regression on the abundance of the parent protein (adjusted phosphopeptides abundance) (**Fig. 1D**). For this purpose, we modified the protein abundance estimation by excluding all peptides corresponding to detected phosphopeptides (**Methods**). In some cases, we were not able to quantify the parent protein after removing phosphorylated peptides, and we obtained 3875, 3471, and 3492 adjusted phosphopeptides in heart, kidney, and liver tissue, respectively.

**Figure 1.**
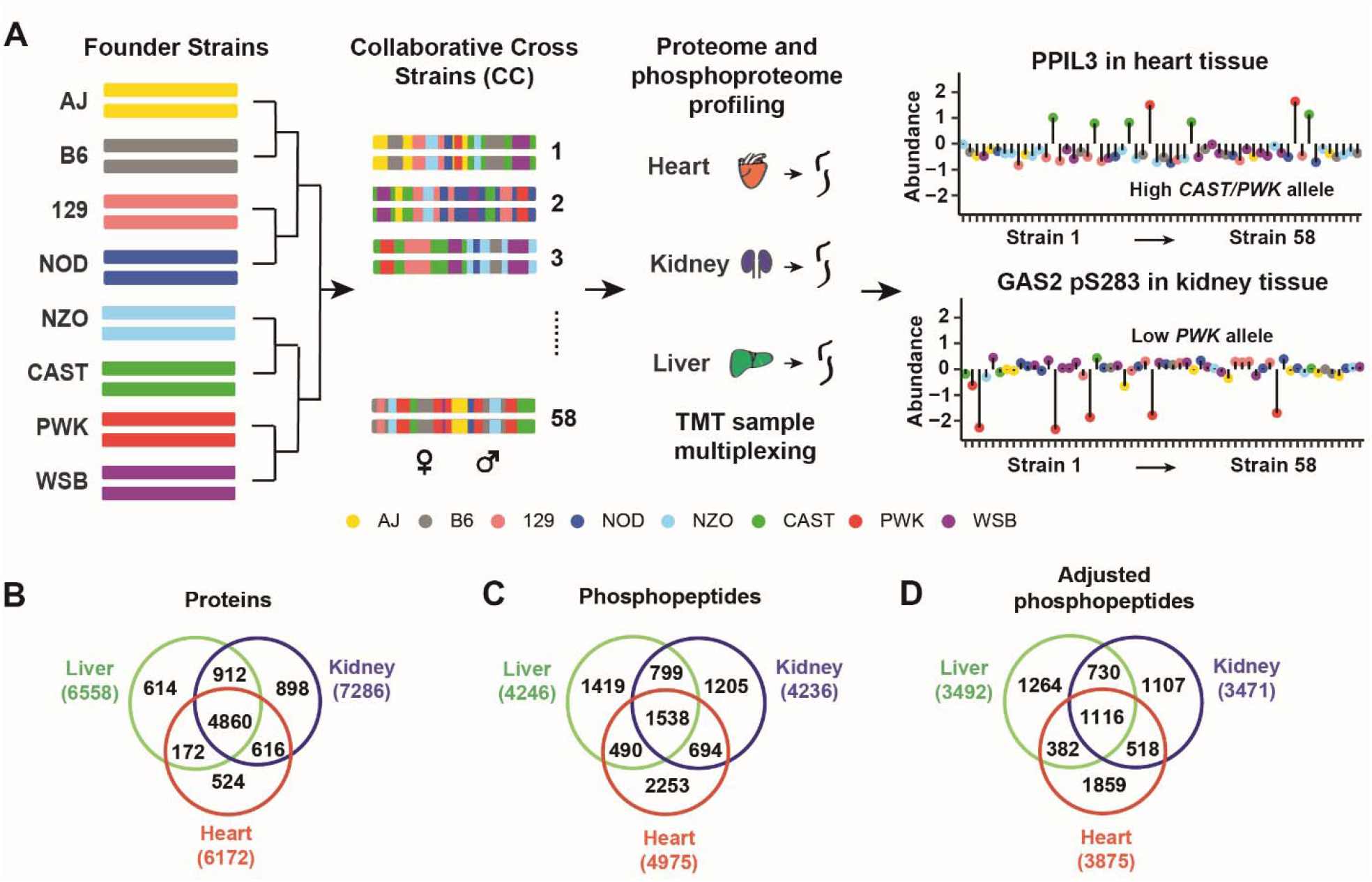
Overview of the proteome and phosphoproteome profiling of three tissues from Collaborative Cross strains using Tandem mass tags (TMT). **(A)** Liver, kidney and heart samples were collected from one male and one female mouse from 58 Collaborative Cross (CC) inbred strains. Samples (116) were multiplexed utilizing TMT sample multiplexing reagents. Proteome and phosphoproteome analyses were collected by mass spectrometry. **(B)** Venn diagrams of the quantified proteins, **(C)** phosphopeptides and (**D**) adjusted phosphopeptides in liver, kidney and heart tissues.

### Sex differences among phosphopeptides

We estimated the effect of sex on abundance of proteins, phosphopeptides, and adjusted phosphopeptides in all three tissues (**Methods**). For heart, we detected significant sex effects (FDR < 0.01) for 323 proteins, 12 phosphopeptides, and 0 adjusted phosphopetides; for kidney, 4,499 proteins, 2,031 phosphopeptides, and 538 adjusted phosphopetides; and for liver, 2,367 proteins, 547 phosphopeptides, and 97 adjusted phosphopetides (**Table S2**). Sex effects are most prevalent in kidney, followed by liver, and there are relatively few in heart **(Fig. 2A**). Standardized sex effects on phosphopeptides and their parent proteins are highly correlated (**Fig. S2A**). After adjustment for parent protein abundance, the magnitude of the sex effects is reduced (**Fig. 2A**), but many remain significant. In addition, we see strong positive correlation of sex effects on phosphopeptides before and after adjustment (**Fig. S2B**). Thus, sex effects on phosphopeptide abundance are determined by sex effects on parent protein abundance and by sex-specific factors that act directly on phosphorylation levels.

**Figure 2.**
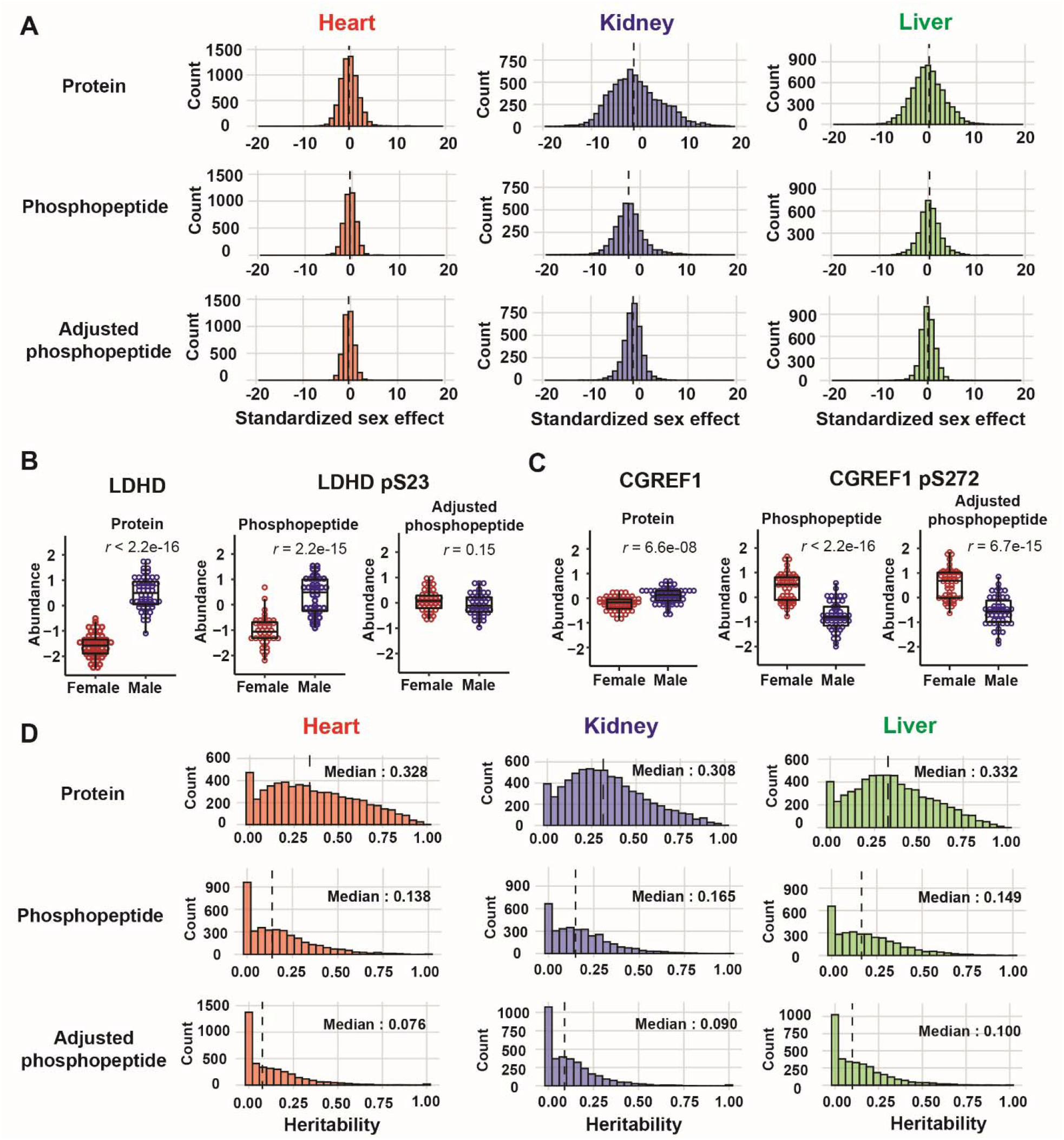
Sex effect and heritability on protein and phosphopepitdes across three tissues. **(A)** Histograms of standardized sex effect (difference/SE) on protein abundance (upper), phosphopeptides (middle), and adjusted phosphopeptides (lower) in heart, kidney and liver tissues. **(B)** Sex difference in the relative abundance (batch corrected log_2_ intensity) of phosphopeptide harboring LDHD pS23 is due to sex effect on its parent protein. **(C)** Sex difference in the relative abundance of phosphopeptide harboring CGREF pS272 is not due to sex effect on its parent protein. **(D)** Histograms of heritability on protein abundance (upper), phosphopeptides (middle) and adjusted phosphopeptides (lower) in heart, kidney and liver tissues. Dashed vertical lines represent the median.

We illustrate how sex can influence phosphopeptide abundance with two examples (**Fig. 2B)**. There is a significant effect of sex on the protein LDHD, which has a higher abundance in males. The phosphopeptide LDHD pS23 also has higher abundance in males, but the adjusted phosphopeptide abundance shows no significant difference between the sexes. We conclude that the sex effect on LDHD pS23 is mediated through the sex effect on the abundance of its parent protein (**Fig. 2B**). The protein CGREF1 has higher abundance in males but CGREF1 pS272 has substantially lower abundance in males. This sex difference in the phosphopeptide persists after adjusting for the parent protein abundance. We conclude that the sex effects on CGREF1 pS272 are mediated by sex-specific processes that act independently of the parent protein abundance (**Fig. 2C**).

### Heritability of phosphopeptides

Heritability is the proportion of phenotypic variation explained by genetic relatedness. It reflects the additive genetic effects on a trait relative to the precision of measurement. We estimated heritability (*h*^*2*^) for the abundance of individual proteins and phosphopeptides in all three tissues. The median heritability across tissues ranged from 0.308 to 0.332 for proteins, from 0.138 to 0.165 for phosphopeptides, and from 0.076 to 0.100 for adjusted phosphopeptides (**Fig. 2D; Table S3**). Protein heritability was substantially higher than phosphopeptide heritability (**Fig. S2C**), which at least in part, reflects the higher precision of protein quantification that combines measurements across multiple peptides. The adjusted phosphopeptides are generally less heritable than the phosphopeptides (**Fig. S2D**), indicating that a dominant component of phosphopeptide heritability is mediated through genetic effects on the parent protein. Nonetheless, there are many adjusted phosphopeptides with non-zero heritability, indicating that genetic factors can directly influence phosphorylation levels.

### Genetic mapping of proteins and phosphopeptides

We mapped pQTL, phQTL and adj_phQTL in all three tissues. We computed a genome-wide adjusted p-value for each trait and then applied a false discovery rate adjustment (FDR < 0.1) to account for the number of proteins or peptides (**Methods**). We identified 1,608, 1,801, and 1,609 pQTL (**Fig. 3A**); 211, 251, and 275 phQTL (**Fig. 3B**); and 40, 58, and 41 adj-phQTL (**Fig. 3B**) in heart, kidney, and liver tissue, respectively (**Table S4**). We defined local QTL as being located within 10 Mbp of the midpoint of the protein-coding gene, all others are distant QTL. Mapping resolution of the CC panel is not uniform across the genome and we noted several instances where QTL classified as distant were cleary local, based on the local LD structure. We see greater sharing across tissues for local pQTL (41% are present in at least two tissues) compared to distant pQTL (11% are present in at least two tissues) (**Fig. S3A**). This is consistent with previous studies on multi-tissue gene expression QTL (eQTL) [19, 39]. The proportion of phQTL shared across tissues is lower, with only five local and one distant phQTL found in all three tissues and 10.5% of all phQTL present across two or more tissues (**Fig. S3B**). The sharing of adj-phQTL is lower still, with only one local (EIF3B pS90; **Figs. 3C, S3C**) and one distant (ATP5A1 pS53) site found across all three tissues and only 10.5% of all adj-phQTL present in two or more tissues. The majority of adj-phQTL have a corresponding phQTL (81.8% of all adj-phQTL) (**Figs. 3B and S3D**). The lower proportion of sharing across tissues for phQTL and adj-phQTL could be due to tissue-specificity of phosphorylation but we cannot rule out reduced mapping power for individual peptides relative to proteins.

**Figure 3.**
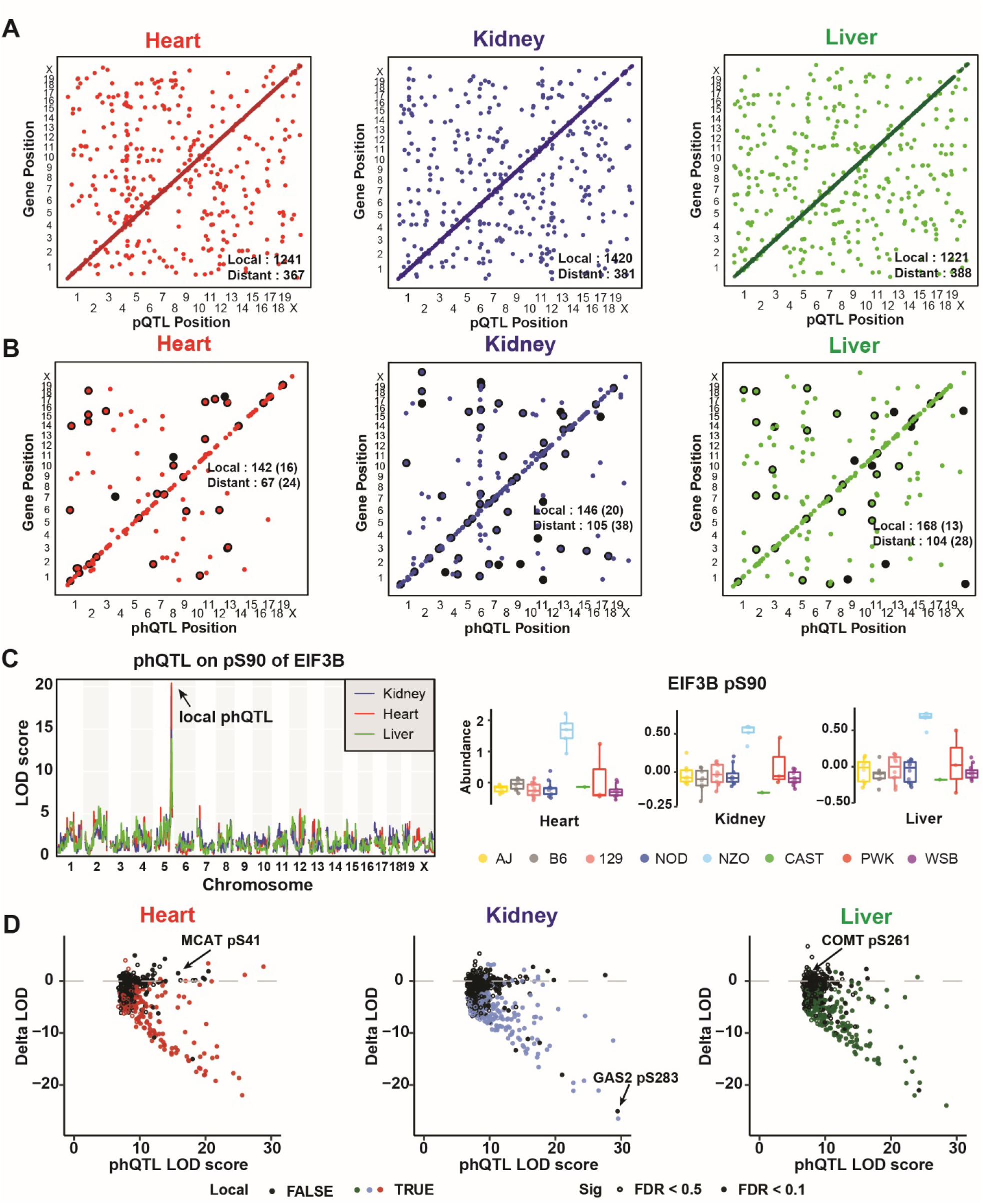
pQTL and phQTL mapping from CC strains in heart, kidney and liver tissues. Stringently detected (FDR < 0.1) **(A)** pQTL, **(B)** phQTL and adjusted phQTL in heart (left), liver (middle) and kidney (right) tissues. QTL are plotted by the genomic positions of proteins against QTL coordinates. Adjusted phQTL were highlighted in black. **(C)** Adjusted phQTL identified on EIF3B pS90 co-mapped in all three tissues. Relative abundances (batch corrected log_2_ intensity) of EIF3B pS90 in each tissue were grouped based on founder local haplotypes. **(D)** LOD scores of local and distant phQTL (FDR < 0.1 or 0.5) changed after adjusting for their parent protein abundances in heart, kidney, and liver tissues. MCAT pS41, GAS2 pS283 and COMT pS261 were labelled.

To determine how much of the genetic contribution to phQTL is mediated through abundance of their parent proteins, we first looked at the correlation of allele effects at concordant pQTL-phQTL pairs. We observed high positive correlations for most pairs, consistent with shared genetic effects (**Fig. S3E**). We then calculated the difference of the LOD score for each phQTL before and after adjusting for parent protein abundance. If the LOD score drops after adjustment, this indicates that the phQTL is mediated, at least in part, through variation in the abundance of the parent protein. The phQTL with the greatest reduction in LOD score (Delta LOD percentage < - 50%, FDR < 0.1) were primarily local phQTL (89.5%-95.7% across tissues), although a few distant phQTL (N = 21) showed a similar reduction in LOD score (**Fig. 3D, Table S5**). We looked at a larger set of phQTL using a less stringent multiple testing correction (FDR < 0.5) and saw the same pattern. We conclude that local genetic effects on phosphopeptide abundance are often mediated through parent protein abundance. However, a substantial number of phQTL, especially those that are distant from the coding gene, show little or no drop in LOD score after adjustment, indicating that these phQTL are responding to genetics effects independent of their parent protein abundance. The drop in LOD scores for many phQTL falls somewhere between these extremes, indicating that they are influenced by parent protein abundance and by independent mechanisms.

We note that the genetic effects on phosphopeptides can be modified by sex. Our experimental design, with one male and one female mouse from each CC strain, is well suited for mapping QTL with genetic effects that differ between the sexes, which we refer to as sex-interactive QTL. We mapped 2, 43, and 5 sex-interactive pQTL (FDR < 0.1) in heart, kidney, and liver, respectively (**Fig. S3F**). We identified 4 sex-interactive phQTL in kidney (3 local and one distant). We found no sex-interactive phQTL in heart or liver and no sex-interactive adj-phQTL in any tissue. The local sex-interactive phQTL for HAO2 pS171 illustrates how sex and genetic variation can simultaneously affect protein and phosphoprotein abundance (**Fig. S3G**). Female mice generally have higher phosphorylation of HAO2 pS171 relative to their male counterparts, but the magnitude of the sex effect is amplified for mice with the CAST allele at this QTL.

### Distant phQTL effects are mediated through kinases, phosphatases, and cytokines

Phosphopeptide abundance can be driven by abundance of the parent protein (Mechanism 1), and by factors that affect phosphorylation levels independently of protein abundance (Mechanism 2; **Fig. 4A**). We set out to quantify the relative contributions of these two mechanisms and to identify candidate mediators of Mechanism 2, which we expected to be enriched for kinases, phosphatases, and upstream regulators of protein phosphorylation.

**Figure 4.**
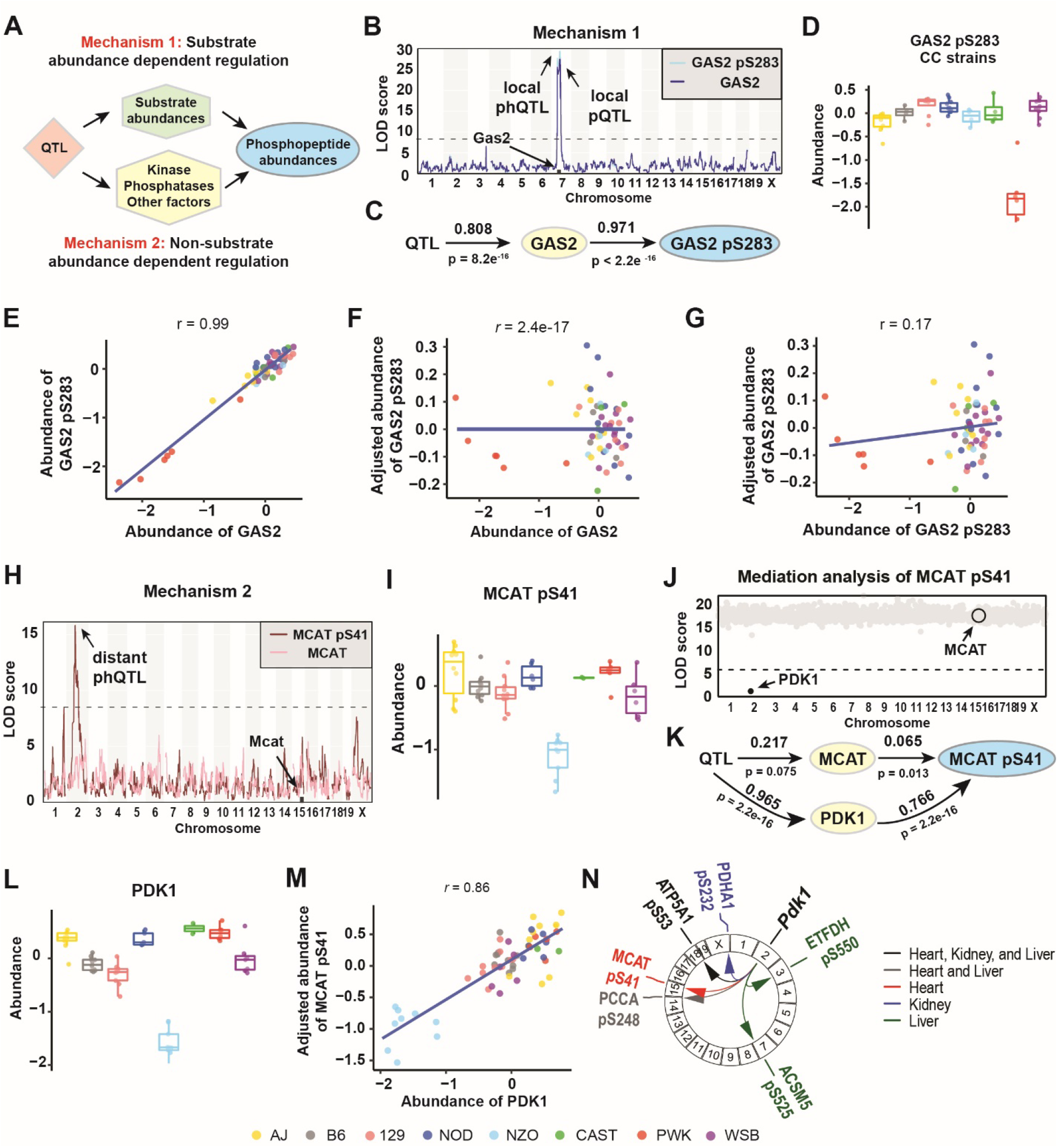
Phosphopeptide abundance can be regulated by substrate abundance dependent or non-substrate abundance dependent mechanisms. **(A)** Diagram showing how the genetic effect resulting in phQTL detection may be regulated by either parent protein abundance (batch corrected log_2_ intensity) changes (Mechanism 1) or by phosphorylation stoichiometry (Mechanism 2) or both. **(B)** Genome scans for GAS2 and GAS2 pS283 in kidney tissue. **(C)** Path diagram of GAS2 pS283 abundance regulation in kidney tissue. **(D)** The *PWK* allele of the GAS2 pS283 phQTL drove low phosphopeptide abundance in kidney tissue. Data were categorized based on the founder haplotye at the identified pQTL. **(E)** Abundances of overall GAS2 and GAS2 pS283 were highly correlated (*r* = 0.99). Points are colored based on founder haplotype at *Gas2*. **(F)** Overall abundance of GAS2 and adjusted abundance (residual from regression of batch corrected log_2_ intensity) of GAS2 pS283 were not correlated (*r* = 2.4e-17). Points are colored based on founder haplotype at *Gas2*. **(G)** Abundance of GAS2 pS283 and adjusted abundance of GAS2 pS283 were not correlated (*r* = 0.02). Points are colored based on founder haplotype at *Gas2*. **(H)** Genome scans for MCAT and MCAT pS41 in heart tissue. **(I)** *NZO* alleles at *Pkd1* drove the low abundances of MCAT pS41 in heart tissue. Colors denote the founder haplotype of additive allele effects at the identified pQTL of MCAT pS41. **(J)** Mediation analysis identified PDK1 expression as the mediator of MCAT pS41 abundances. Each gray dot is a mediation score representing the MCAT pQTL LOD score conditioned on a protein as candidate mediator. **(K)** Path diagram of MCAT pS41 abundance regulation in heart tissue. **(L)** *NZO* alleles at *Pkd1* drove the low abundances of PDK1 in heart tissue. Colors denote the founder haplotype of additive allele effects at the identified pQTL of MCAT pS41. **(M)** The adjusted abundances of MCAT pS41 and PDK1 were highly correlated (*r* = 0.86) in heart tissue. **(N)** Mediation analysis identified PDK1 as the mediator of several phQTL in heart, kidney and liver tissue, respectively.

We observed that many phQTL have a corresponding pQTL but no correpsonding adj-phQTL, *i*.*e*., the LOD score drops when the phosphopeptide is adjusted for the parent protein abundance (**Fig. 3D**). The genetic effects at these phQTL are mediated by Mechanism 1. For example, a pQTL on chromosome 7 at 64Mb explains 81% of variation in the abundance of the protein GAS2 in kidney (p = 8.2e-16) (**Fig. 4B-C**). The abundance of GAS2 is low in animals with the PWK allele at this locus (**Fig. 4D**). The abundance of GAS2 pS283 is highly correlated with its parent protein’s abundance (r = 0.99) (**Fig. 4E**). After adjusting for GAS2 abundance, GAS2 pS283 is no longer associated with the genotype at the phQTL locus (**Fig. 4G**). Additional examples of phQTL that are mediated through parent proteins include TPMT pS34 (**Fig. S4A-D**) and MTX3 pS284 (**Fig. S4E-H**). Common features of phQTL consistent with Mechanism 1 are a strong local pQTL for the parent protein and strong correlation between the parent protein and the phosphopeptide.

We observed 74 distant phQTL that had no corresponding pQTL, and after adjusting for parent protein abundance, the adj-phQTL remained significant (**Fig. 3D**). The genetic effects at these QTL are mediated primarily by Mechanism 2. For example, MCAT pS41 in heart (**Fig. S5**) has distant phQTL and adj-phQTL on chromosome 2 at 71.8Mb (**Fig. 4H**). The abundance of MCAT pS41 is low when this Chr 2 locus carries an NZO allele (**Fig. 4I**). To identify the gene candidates responsible for this effect, we applied mediation analysis to evaluate the transcripts and proteins in the phQTL region on Chr 2 (**Methods**). The strongest mediation signature was found for PDK1, *pyruvate dehydrogenase kinase 1* (**Fig. 4J-K**). The transcript abundance of *Pdk1* was also identified as a mediator. PDK1 has a local pQTL with low expression in mice with an NZO allele (**Fig. 4L**), and PDK1 abundance is tightly correlated with MCAT pS41 (**Fig. 4M**). The pQTL of PDK1 explains 97% (p < 2.2e-16) of the variation in PDK1 abundance and 77% (p < 2.2e-16) of variation in MCAT pS41 abundance. The effect of the pQTL on MCAT was not significant and the effect of MCAT abundance on MCAT pS41 was significant but weak (**Fig. 4K**), confirming that the phQTL on MCAT pS41 was not mediated through MCAT abundance but is primarily driven by PDK1 abundance. Across all three tissues, we found a total of 9 distant phQTL (on 6 different proteins) that map to the *Pdk1* locus on Chr 2 and are mediated by PDK1, including the confirmed substrate of PDK1, pyruvate dehydrogenase E1 component subunit alpha, PDHA1 [40] (**Fig. 4N**). The phQTL at Chr 2 for ATP5A1 pS53 is found in all three tissues, and the phQTL for PCCA pS248 is found in heart and liver tissues. These results indicate that PDK1 is the upstream kinase of these phosphorylation sites.

We found 45 examples of phosphopeptides whose abundance is influenced by both mechanisms 1 and 2 to different degrees and with genetic associations that are local, distant, or both. For example, the protein COMT in liver has a local pQTL on Chr 16 at 18Mb, and COMT pS261 has a distant adj-phQTL on Chr 13 at 54.8Mb (**Fig. 5A**). The local pQTL drives higher expression of COMT in the presence of a CAST allele (**Fig. 5B**). After adjusting for COMT abundance, the adjusted phosphopeptide shows high abundance in the presence of a WSB allele at the distant phQTL (**Fig. 5C**). Mediation analysis of the Chr 13 QTL identified the transcript of *Cdc14b* as a candidate mediator of phosphorylation (**Fig. 5D**). We note that CDC14B was not quantified in the proteomics analysis. The distant adj-phQTL for COMT pS261 co-maps with a local eQTL for *Cdc14b* on Chr 16 and exhibit mirrored allele effects, *i*.*e*., the WSB allele confers low expression of *Cdc14b* but high abundance of COMT pS261, resulting in negative correlation between COMT pS261 and *Cdc14b* mRNA abundance. Regressing out the effect of COMT protein abundance on the abundance of the COMT pS261 phosphopeptide improves this correlation between *Cdc14b* mRNA and COMT pS261, which confirms that abundance of COMT pS261 phosphopeptide is regulated by both its parent protein abundance and the transcript abundance of *Cdc14b* (**Figs. 5E-G**). The Chr 13 QTL explains 63% (p = 5.2e-9) of variation in COMT and in turn, COMT explains 48% (p = 2.6e-9) of variation in COMT pS261. The QTL on Chr 13 explains 51% (p = 4.2e-6) of variation in *Cdc14b*, which in turn explains 38% (p = 2.8e-7) of variation in COMT pS261.

**Figure 5.**
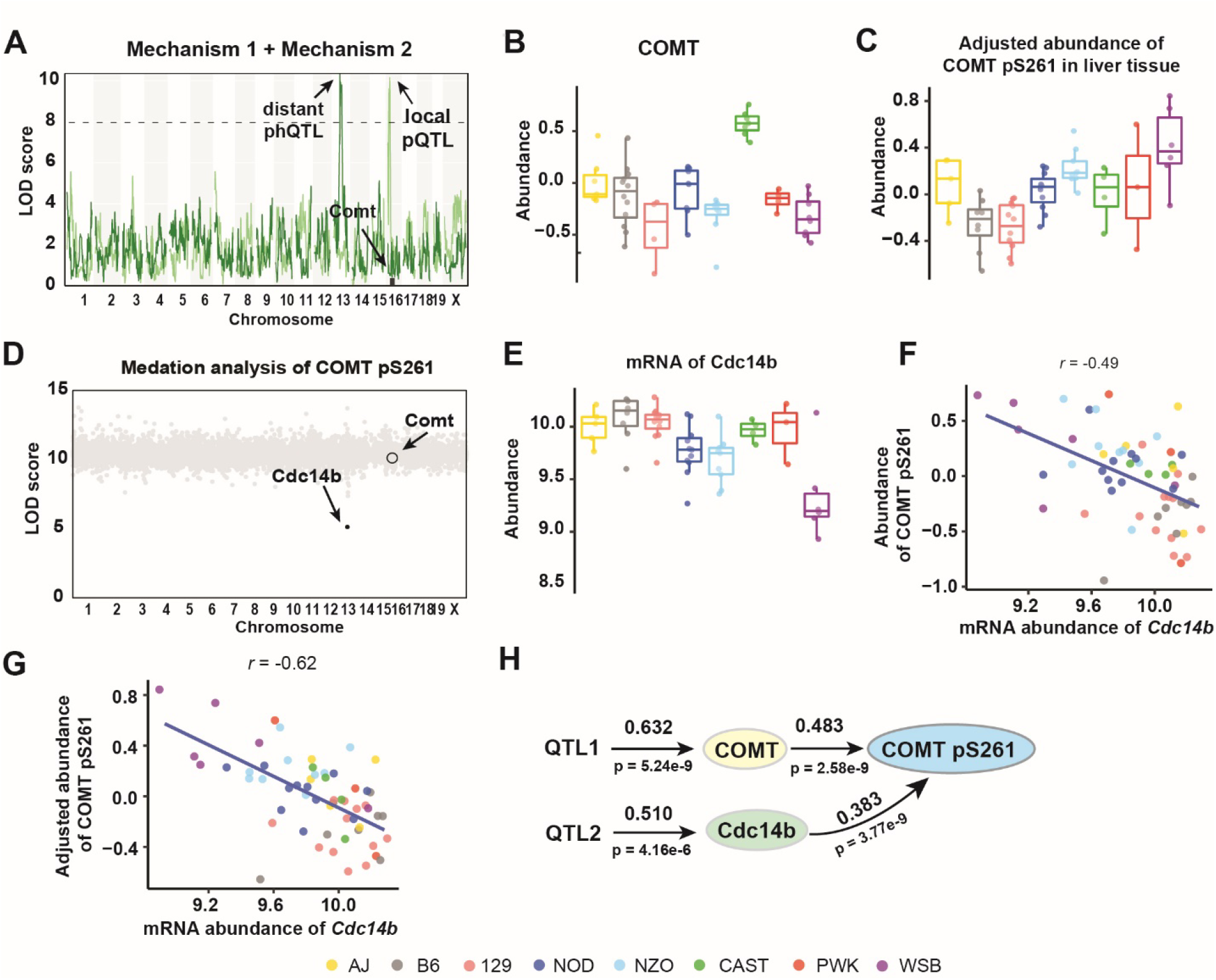
Phosphopeptide abundance can be regulated by both substrate abundance dependent and non-substrate abundance dependent mechanisms. **(A)** COMT pQTL and phQTL for COMT were mapped to different loci in liver tissue. **(B)** A local *CAST* allele at *Comt* drove high abundance of COMT in liver tissue. **(C)** Adjusted abundance of COMT pS261 categorized according to founder haplotype at *Cdc14b*. **(D)** Mediation analysis using transcriptomics data identified Cdc14b as the mediator of a phQTL for COMT pS261. Each gray dot is a mediation score representing the COMT pS261 phQTL LOD score conditioned on a transcript as candidate mediator. **(E)** Abundance of *Cdc14b* transcripts pS261 categorized according to founder haplotype at *Cdc14b*. The abundance of COMT pS261 is less correlated with *Cdc14b* transcripts before adjustment (*r* = -0.49) **(F)** compared to after adjustment **(G)** (*r* = - 0.62). **(H)** Path diagram of COMT pS261 abundance regulation in liver tissue.

A second example of complex regulation, LMNA pS394 was also found to be mediated by *Cdc14b* in heart (**Fig. S6 I-L**). *Cdc14b* is a dual specificity protein phosphatase known to be involved in DNA damage response [41] and cell cycle regulation [42], and based on this genetic data is the likely upstream phosphatase acting on COMT pS261 and LMNA pS394 in liver and heart, respectively. Additional examples with complex genetic regulation include PDLIM4 pS119 and NGEF pS606, both found in heart (**Fig. S6**). Genetic effects on PDLIM4 pS119 were mediated through PDLIM4 abundance and *Il15* transcript expression (**Fig. S6A-F**). *Il15* is a cytokine, and signaling through *Il15* results in kinase SYK activation to stimulate cell proliferation [43]. Allele effects of the phQTL (PDLIM4 pS119), the eQTL (*Il15*), and mediation analysis are all consistent with higher levels of *Il15* leading to higher levels of PDLIM4 pS119. For NGEF pS606, we found that its distant phQTL was mediated through the transcript abundance of *Prkca*, protein kinase C, alpha (**Fig. S6G-L**).

In summary, we found that most local phQTL have a corresponding pQTL and are primarily driven by their parent protein abundance (mechanism 1), while distant phQTL with adj-phQTL are primarily driven by factors that are independent of the parent protein abundance (mechanism 2). We identified many examples of regulation of phosphopeptides by both mechanisms 1 and 2 (**Table S5**). These include 6 kinases (*Pdk1, Mapkapk3, Nme6, Plk2, Prkca, Sbk3*), 3 phosphatases (*Cdc14a, Cdc14b*, Pxylp1) and additional genes that are known to be involved in cell signaling transduction and affect protein phosphorylation, including *Il15, Negr1*, and *Stat6*.

### Regulation of phosphorylation sites within a protein

We next asked whether phosphopeptides that co-occur on the same protein were coordinately regulated. We identified 1151, 1148, and 1093 proteins with two or more phosphopeptides quantified in heart, kidney, and liver tissues, respectively (**Fig. S1A**). To determine whether phosphorylation sites on the same protein were potentially co-regulated, we looked at the correlation of the abundances of phosphopeptides from the same protein. (**Fig. S7A-B**). In each tissue, the median correlation of phosphopeptides decreased but remained significant after adjustment based on their parent proteins, indicating that phosphopeptides from the same protein can be co-regulated independently of parent protein abundance. For example, we quantified 7 phosphopeptides from EGFR in liver tissue with correlations among the adjusted phosphopeptides ranging from -0.049 and 0.598 (**Fig. 6A**). While only one of these sites had a significant adj-phQTL (pS1044, Chr 9 at 107Mb), two sites had sub-threshold adj-phQTL with allele effects that are consistent with a shared adj-phQTL, suggesting that phosphorylation sites on EGFR can be co-regulated.

**Figure 6.**
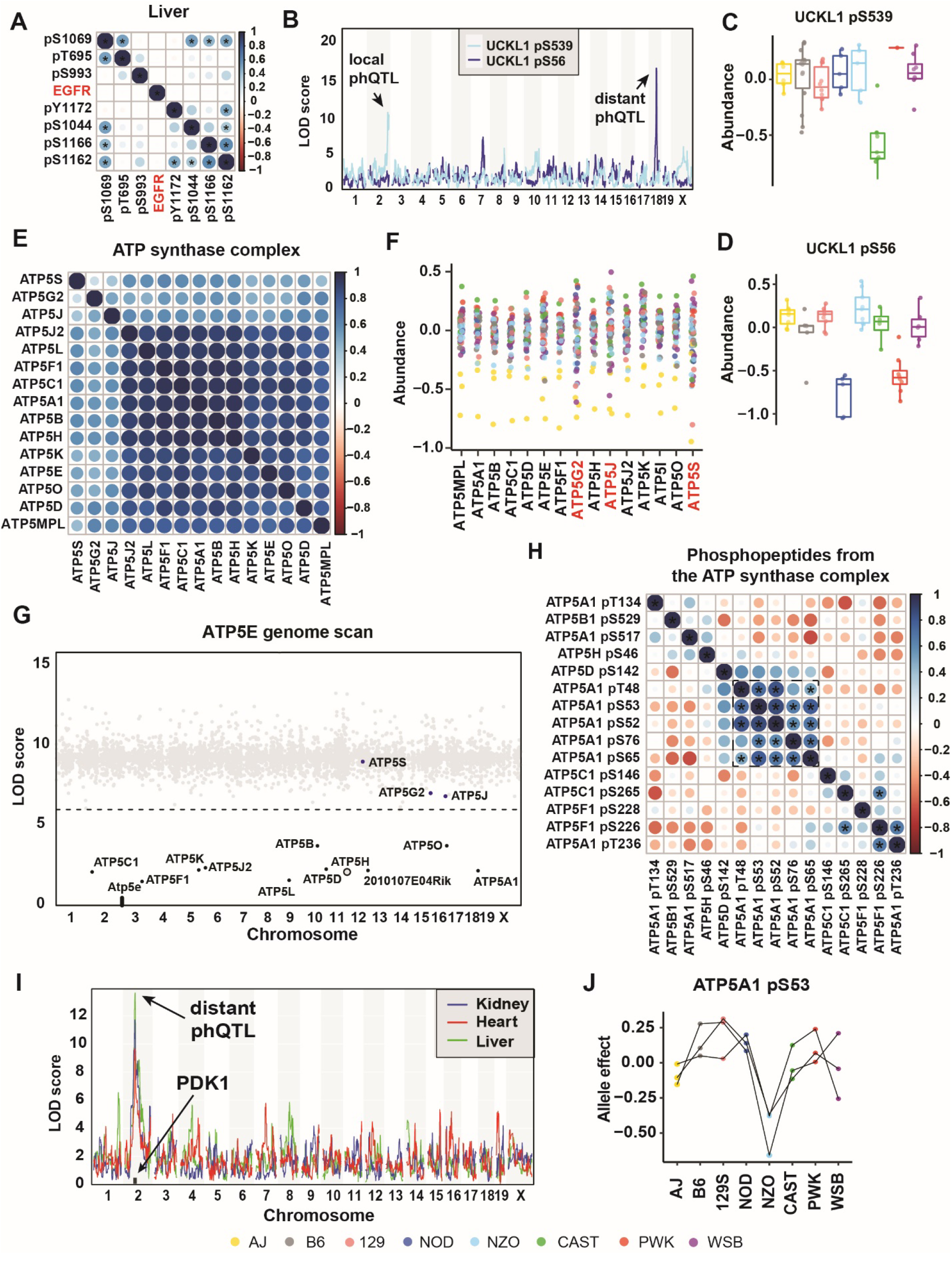
Phosphorylation sites on one protein can be regulated coordinated and not coordinated. **(A)** Heatmap of Pearson correlations of abundances of phosphopeptides from parent protein ABLIM1. **(B)** Genome scans of pS539 and pS56 on UCKL1 in kidney tissue. **(C)** A local CAST allele at *Uckl1* drove low abundance of UCKL1 pS539 in kidney tissue. **(D)** Distant NOD *and* PWK allele on chromosome 18 drove low abundance of UCKL1 pS56 in kidney tissue. **(E)** Heatmap of Pearson Correlations among all proteins quantified in ATP synthase complex in heart tissue. **(F)** The *AJ* allele at *Atp5h* drove low abundance of the entire ATP synthase complex in heart tissue. All quantified ATP synthase complex subunits have low protein abundance in CC032, CC033 and CC044 strains, which possess the *AJ* allele, in heart data. **(G)** Mediation analysis using proteomics data identified ATP5H as the mediator of a phQTL for ATP5E. Each gray dot is a mediation score representing the ATP5E pQTL LOD score conditioned on a protein as candidate mediator. ATP5H was detected as the strongest mediator of the ATP5E distal pQTL in heart tissue. All ATP synthase complex subunits have mediation z-scores < -8 and were highlighted in black. Other quantified ATP synthase complex subunits, ATP5S, ATP5G2 and ATP5J, were highlighted in blue. Horizontal dashed line at LOD of 6 was included for reference. **(H)** Heatmap of Pearson Correlations among all phosphorylation events quantified from the ATP synthase complex in heart tissue. The correlations among the five sites from ATP5A1 are highlighted by a dashed square. Correlation with FDR < 0.01 were highlighted using stars. **(I)** Genome scans of ATP5A1 pS53 in the three tissues, revealing co-mapping phQTL in all the three tissues. **(J)** Allele effects of ATP5A1 pS53 phQTL were highly correlated in the three tissues.

Phosphorylation sites on one protein can be regulated differently. For example, abundances of UCKL1 pS56 and UCKL1 pS539 are not correlated (*r* = 0.020), and genetic mapping identifies a local phQTL on chr 2 for UCKL1 pS539 and a distant phQTL on chr 18 for UCKL1 pS56 (**Fig. 6B**). The CAST allele drives the low abundance of UCKL1 pS539 (**Fig. 6C**), and NOD and PWK alleles drive the low abundance of UCKL1 pS56 (**Fig. 6D**), presenting an example of phosphorylation sites on one protein that are regulated by distinct mechanisms. An adj-phQTL was identified for UCKL1 pS56 but not for UCKL1 pS539. We conclude that the local phQTL of UCKL1 pS539 was mediated through protein abundance (Mechanism 1), whereas the phQTL of UCKL1 pS56 was independent of protein abundance regulation (Mechanism 2).

### Genetic regulation of the ATP Synthase Complex

We conclude with two examples that illustrate how these data can be used as a resource to dissect the genetic regulation of protein and phosphopeptide abundance. The first example is the ATP synthase complex, which is localized to the inner mitochondrial membrane where it converts ADP to ATP as the final step of oxidative phosphorylation [44]. In heart tissue, we quantified 15 subunits of the complex and detected 15 phosphopeptides. The complex is present in kidney and liver as well, but fewer proteins and phosphopeptides were detected in these tissues.

Previously, we demonstrated that proteins that form complexes are often co-regulated [24, 25]. The abundance of subunits from the ATP synthase complex are tightly correlated (median correlation *r* = 0.83) (**Fig. 6E**). In heart tissue, several subunits share a significant co-mapping distant pQTL on Chr 11 at 96Mb, which is the location of the *Atp5h* gene. Mediation analysis of the distant pQTL identified ATP5H as the mediator of complex-wide protein abundance (**Table S5**). The A/J allele at *Atp5h* is associated with low complex-wide abundance, consistent with stoichiometric regulation of the complex by the lowest expressed subunit **(Fig. 6F-G**) [24, 25].

We looked at phosphorylation sites across the complex in heart tissue. The abundance of phosphopeptides from the ATP synthase complex are less tightly correlated (median correlation *r* = 0.049) compared to the proteins (**Figs. 6E, 6H**). Similar results were seen in liver and kidney (**Fig. S7C-F**). Among the 15 phosphorylation sites detected, a cluster of sites in ATP5A1 including pS53 are highly correlated, and share a suggestive (FDR < 0.5) genetic association with the *Pdk1* locus. ATP5A1 pS53, which is quantified in all three tissues, has a distant adj-phQTL on Chr 2 at 73 Mb that is mediated by PDK1 (**Fig. 6I, S7C**), and has low levels of phosphorylation associated with the NZO allele at this locus (**Fig. 6J**). We also identified two significant (FDR < 0.01) correlations between sites in different subunits: ATP5F1 pS226 and ATP5A1 pT236, and ATPF1 pS226 and ATP5C1 pS265, suggesting possible coordination of phosphorylation activity across subunits within the Atp5 synthase complex.

### Genetic regulation of propionyl-CoA carboxylase

PCCA and PCCB together make up the biotin-dependent propionyl-CoA carboxylase (PCC), a mitochondrial enzyme involved in the catabolism of odd chain fatty acids and branched-chain amino acids [45, 46]. A single phosphorylation site pS248 on PCCA was detected in all three tissues, whereas no phosphopeptides were found from PCCB. The site PCCA pS248 had a significant distant adj-phQTL on Chr 2 at 72Mb in both heart and liver. There is a suggestive distant adj-phQTL at the same locus in kidney (LOD = 6.7). In all three tissues, the Chr 2 QTL had a low NZO allele (**Fig. 7A-B**) and was mediated through PDK1 (**Fig. 7C**).

**Figure 7.**
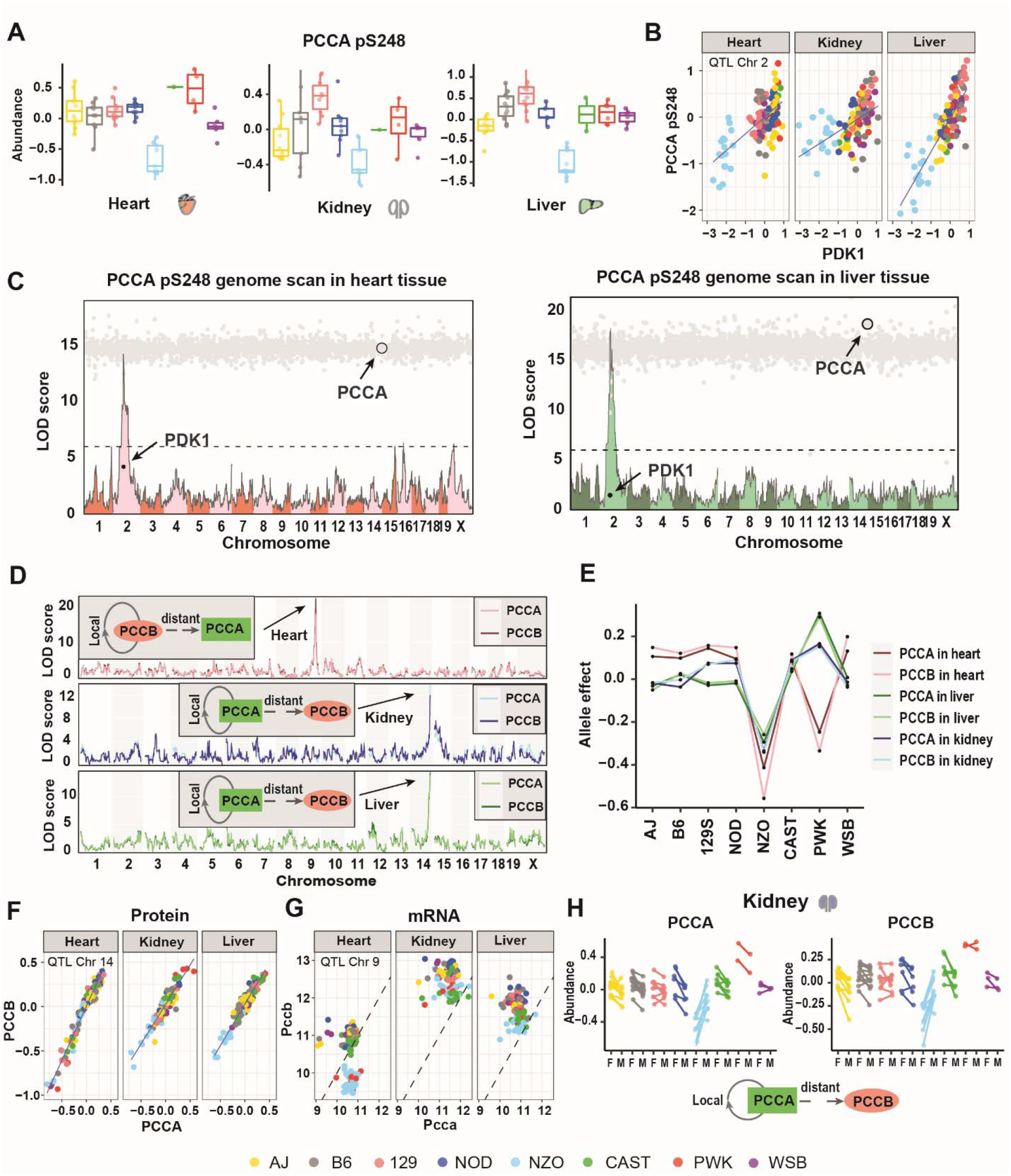
Genetic regulation of PCCA and PCCB across three tissues. **(A)** Co-mapping distant phQTL of PCCA pS248 was identified in liver, heart but not kidney tissue. NZO allele drove the low level of this phosphorylation event. Data were categorized based on the founder haplotye at the identified phQTL. **(B)** Abundances of PDK1 and PCCA pS248 were highly correlated in heart and liver but not in kidney tissue. Abundance of PDK1 and PCCA pS248 in each individual sample (116) were categorized based on the haplotye of the phQTL on PCCA pS248 in heart and liver tissues on Chromosome 2. **(C)** Genome scans for PCCA pS248 are overlayed with mediation scores in heart and liver tissues. Each gray dot is a mediation score representing the PCCA pS248 phQTL LOD score conditioned on a protein as candidate mediator. **(D)** Genome scans of PCCA and PCCB in all the three tissues. Local-pQTL for PCCB and distant-pQTL for PCCA co-mapped to the same locus in heart tissue. PCCB was identified as the mediator of the PCCA distant-pQTL. Local-pQTL for PCCA and distant-pQTL for PCCB co-mapped to the same locus in liver tissue and kidney tissues. PCCA was identified as the mediator of the PCCB distant-pQTL. **(E)** Allele effects of identified pQTL for PCCA and PCCB in the three tissues. **(F)** Protein abundance of PCCA and PCCB were highly correlated in each tissue. Protein abundance in each individual sample (116) were categorized based on the haplotye of the pQTL on PCCA in kidney and liver tissues on Chromosome 14. **(G)** The transcript level of *Pccb* is distinctly higher than the mRNA level of *Pcca* in kidney and liver tissues but not in heart tissue. mRNA abundance of *Pcca* and *Pccb* in each individual sample (116) were categorized based on the haplotye of the pQTL on PCCA in heart tissue on Chromosome 9. **(H)** Sex interactive local pQTL on PCCA and sex-interactive distant pQTL on PCCB co-mapped to the same locus in the kidney tissue, characterized by a distinct NZO effect (NZO males with greater abundances than NZO females). PCCA was identified as the mediator of PCCB sex interactive distant pQTL. Points are colored by founder haplotype at sex-interactive phQTL. Males and females from the same CC strain were connected by a line.

We also identified a local pQTL on PCCB and PCCA has a co-mapping distant pQTL (Chr 9 at 100Mb) that is mediated by PCCB in heart tissue. Low expression of PCCA and PCCB is associated with the NZO and PWK alleles at the Chr 9 QTL. In kidney and liver, PCCA instead maps with the a local pQTL and PCCB has a distant pQTL (Chr 14 at 123Mb), mediated by PCCA (**Fig. 7D**). Low expression at the Chr 14 QTL is associated with the NZO allele. The protein abundances of PCCA and PCCB are tightly correlated in both tissues (r = 0.935 in kidney to r = 0.975 in heart, p < 2.2e-16; **Fig. 7E-F**). We hypothesized that the switching across tissues of the local and distant QTL for protein abundances was due to tissue-specific changes in stoichiometric regulation. This is confirmed by looking at the mRNA level at these genes (**Fig. 7G**). In kidney and liver, *Pcca* mRNA has lower abundance and the Chr 14 QTL (local to *Pcca*) is the common driver of PCCA and PCCB protein abundance. In heart, when NZO or PWK alleles are present at the Chr 9 locus (local to *Pccb*), the mRNA level of *Pccb* is lower than *Pcca*, and PCCB becomes the driver of protein abundances. This is consistent with stoichiometric regulation in which the gene with lowest mRNA expression becomes the genetic driver of protein complex abundance [24]. We also note that the Chr 14 pQTL for PCCA (local) and PCCB (distant) in the kidney is sex-specific, with low expression in the presence of the *NZO* allele being most pronounced in females (**Fig. 7H**).

## Discussion

We quantified transcripts, proteins, and phosphorylated peptides across three tissues in a genetically diverse mouse population. Examining the adjusted phosphopeptides, we demonstrated that phosphorylation levels are heritable and can differ between sexes. We mapped pQTL, phQTL, and adj-phQTL and describe two distinct mechanisms for genetic regulation of phQTL. A large proportion of phQTL are mediated through protein abundance. Other phQTL remain significant after accounting for the effects of the parent protein abundance on phosphopeptide abundance (adj-phQTL) suggesting that genetic factors are likely affecting the levels of site-specific phosphorylation. We applied mediation analysis to identify proteins or transcripts that are candidate causal intermediates underlying distant adj-phQTL. These mediators included kinases, phosphatases, and upstream regulators involved in the phosphorylation process. We highlighted the most significant mediation effects above. However, there are many more examples of plausible mediation that can be mined from these data (Table S5), providing experimentally testable hypotheses about the molecular interactions that mediate site-specific phosphorylation.

We identified PDK1 abundance as the mediator of nine adj-phQTL across three tissues. All nine phosphorylation sites were found in parent proteins involved in the respiratory chain, including sites on APT5A1 and PCCA with adj-phQTL that are shared across all three tissues. Looking at genetic variants at the *Pdk1* locus, we identified a 2bp insertion (Chr 2: 71874272 – 71874274, GRCm38) in the promoter that occurs only in NZO mice and may drive the low expression of PDK1, leading to lower kinase activity and ultimately lower levels of phosphorylation on key proteins involved in respiratory chain metabolism. These findings are particularly interesting because the NZO mouse is a well-studied polygenic model for human metabolic syndrome [47, 48]. QTL mapping in the NZO mouse, has identified *Tbc1d1* [49], *Zfp69* [50] and *Lepr* [51], as genes contributing to type 2 diabetes. Here, we identify a potential role for aberrant protein phosphorylation due to low expression of PDK1 that may further contribute to metabolic disease phenotypes characteristic of the NZO mouse.

While investigating the protein complex formed by PCCA and PCCB, besides the adj-phQTL identified on pS248 on PCCA, we identified two additional pQTL, one local to PCCA and the other local to PCCB, both with low expression of the NZO allele. In heart tissue, the C to T (100,982,310bp) and G to A (100,987,863bp) mutations specific to the NZO and PWK alleles potentially affect transcription and lead to the lower transcript level and protein level of PCCB (100,864,085bp-100,916,951bp). In kidney and liver tissues, NZO specific mutations in *Pcca* may cause the low abundance of PCCA and PCCB. This example illustrates the complexity and delicacy of the mechanisms of genetic regulation of protein abundance and phosphorylation.

We also recognize some limitations of the current study. We find suggestive evidence for many genetic effects on phosphopetides that did not reach stringent genome-wide and multiple testing adjusted significance criteria. The CC panel is limited and it is not possible to improve mapping power substantially by adding more strains. However, by adding more animals per strain, the precision of protein and peptide quantification can be improved to increase mapping power [23]. There were also instances where we did not detect phosphorylated peptides that must be present, for example, the ATP synthase complex in liver and kidney. Advanced mass spectrometry technology, especially targeted mass spectrometry technology could be developed and used to obtain better coverage and provide a more complete picture of the phosphorylated proteome [52].

This integrative multi-omics analysis in genetically diverse CC strains provides a powerful tool to identify regulators of protein-phosphorylation. Similar approaches could be used in combination with interventions, including mapping modifiers of transgenic models of disease [53]. The multi-omics data generated in this study provides a resource for further exploration. The upstream kinases, phosphatases, or other regulating factors identified here can seed hypotheses and motivate further mechanistic studies in disease models. Moreover, it sets a precident for future studies of regulatory mechanisms for other post translation modifications (PTMs) of proteins, such as methylation and ubiquitination. Coupled with advanced mass spectrometry technology for deeper coverage, we foresee this strategy being used to provide a comprehensive regulatory map of PTMs.

## METHODS

### Mice

We received pairs of young mice from 58 CC strains from the UNC Systems Genetics Core Facility between the summer of 2018 and early 2019. Mice were singly housed upon receipt until eight weeks of age. More information regarding the CC strains can be found at https://csbio.unc.edu/CCstatus/index.py.

### Genotyping, founder haplotype reconstruction, and gene annotation

The genotyping and haplotype reconstruction for the CC mice were previously described [25]. Briefly, the 116 CC mice were genotyped on the Mini Mouse Universal Genotyping Array [54] (MiniMUGA), which includes 11,125 markers. Founder haplotypes were reconstructed using a Hidden Markov Model (HMM), implemented in the qtl2 R package [55], using the “risib8” option for an eight-founder recombinant inbred panel and Genome Reference Consortium Mouse Build 38 (mm10). Heterozygous genotypes were omitted, and haplotype reconstructions are limited to homozygous states, smoothing over a small number of residual heterozygous sites that remain in the CC mice. Ensembl v91 gene and protein annotations were used in the CC and founder strains.

### Sample preparation for proteomics and phosphoproteomics analysis

Proteome sample preparation and data analysis for the CC liver tissue was described previously [25]. We also collected kidney and heart tissues along with liver tissue. Singly housed CC mice had their food removed six hours prior to euthanasia and tissue harvest. Tissues were dissected, weighed, and snap frozen in liquid nitrogen. Pulverized heart and kidney tissue were syringe-lysed in 8 M urea and 200 mM EPPS pH 8.5 with protease inhibitor and phosphatase inhibitor. BCA assay was performed to determine protein concentration of each sample.

Samples were reduced in 5 mM TCEP, alkylated with 10 mM iodoacetamide, and quenched with 15 mM DTT. 100 μg protein was chloroform-methanol precipitated and re-suspended in 100 μL 200 mM EPPS pH 8.5. The proteins were digested by Lys-C at a 1:100 protease-to-peptide ratio overnight at room temperature with gentle shaking. Trypsin was used for further digestion for 6 hours at 37°C at the same ratio with Lys-C. After digestion, 50 μL of each sample were combined in a separate tube and used as the 16^th^ sample in all 8 tandem mass tag (TMT) 16plex, rather than the 11plex used previously for liver tissue. 50 μL of each sample were aliquoted, and 12 μL acetonitrile (ACN) was added into each sample to 30% final volume. 100 μg TMT reagent (126, 127N, 127C, 128N, 128C, 129N, 129C, 130N, 130C, 131N, 131C, 132N, 132C, 133N, 133C, 134N) in 10 μL ACN was added to each sample. After 1 hour of labeling, 1 μL of each sample was combined, desalted, and analyzed using mass-spec. Total intensities were determined in each channel to calculate normalization factors. After quenching using 0.3% hydroxylamine, 16 samples were combined in 1:1 ratio of peptides based on normalization factors.

High-Select Fe-NTA Phosphopeptide Enrichment Kit (Thermo Fisher) was used to enrich the phosphorylated peptides (phosphopeptides) according to the manufacturer’s protocol. Flow through and washes from phosphopeptide enrichment were combined, dried, and fractionated with basic pH reversed phase (BPRP) high performance liquid chromatography (HPLC) as described before. We used an Agilent 1260 pump equipped with a degasser and a single wavelength detector (set at 220 nm). Peptides were subjected to a 50 min linear gradient from 8% to 40% acetonitrile in 10 mM ammonium bicarbonate pH 8 at a flow rate of 0.6 mL/min over an Agilent 300Extend C18 column (3.5 μm particles, 4.6 mm ID and 250 mm in length). The peptide mixture was fractionated into a total of 96 fractions which were consolidated into 24. Twelve fractions were desalted and analyzed by liquid chromatography-tandem mass spectrometry (LC-MS/MS). Meanwhile, the eluant from the phosphopeptide enrichment was desalted and analyzed by LC-MS/MS.

### Liquid chromatography and tandem mass spectrometry

The method for proteome data collection in liver tissue was described previously [56]. Proteome data in heart and kidney tissues were collected on an Orbitrap Eclipse mass spectrometer coupled to a Proxeon NanoLC-1200 UHPLC. The peptides were separated using a 100 μm capillary column packed with ∼35 cm of Accucore 150 resin (2.6 μm, 150 Å; ThermoFisher Scientific). The mobile phase was 5% acetonitrile, 0.125% formic acid (A) and 95% acetonitrile, 0.125% formic acid (B). For BPRP fractions, the data were collected using a DDA-SPS-MS3 method with online real-time database searching (RTS) [57] [58]. The data were collected using a DDA-SPS-MS3 method. A database that included all entries from an indexed Ensembl mouse database version 90 (downloaded:10/09/2017) was used in RTS. Each fraction was eluted using a 90 min method over a gradient from 6% to 30% B. Peptides were ionized with a spray voltage of 2,500 kV. The instrument method included Orbitrap MS1 scans (resolution of 1.2×10^5^; mass range 400−1600 m/z; automatic gain control (AGC) target 4×10^5^, max injection time of 50 ms and ion trap MS2 scans (CID collision energy of 35%; AGC target 7.5×10^3^; rapid scan mode; max injection time of 50 ms). RTS was enabled and quantitative SPS-MS3 scans (resolution of 50,000; AGC target 2×10^5^; max injection time of 200 ms) were processed through Orbiter real-time database searching. This data acquisition includes high-field asymmetric-waveform ion-mobility spectrometry (FAIMS). The dispersion voltage (DV) for FAIMS was set at 5000V, the compensation voltages (CVs) were set at -40V, -60V, and -80V, and TopSpeed parameter was set at 1 sec per CV [59].

Mass spectrometric data for phosphopeptides fractions in liver tissue were collected on an Orbitrap Lumos mass spectrometer. Mass spectrometric data were collected in HCD and CID modes. Each fraction was eluted using a 180 min method over a gradient from 6% to 30% B. Peptides were ionized with a spray voltage of 2,600 kV. The instrument method included Orbitrap MS1 scans (resolution of 1.2 ×10^5^; mass range 400−1400 m/z; automatic gain control (AGC) target 1×10^6^, max injection time of 50 ms. The 10 most intense MS1 ions were selected for MS2 analysis. Following acquisition of each MS2 spectrum, a synchronous-precursor-selection (SPS) MS3 scan was collected on the Top 10 most intense ions in the MS2 spectrum. The isolation width was set at 0.7 Da and isolated precursors were fragmented using two methods. In the first method, we used collision induced dissociation (CID) at a normalized collision energy (NCE) of 35% with MultiStage Activation (MSA), and in the second method we used higher energy collision-induced dissociation (HCD) at a normalized collision energy (NCE) of 33%. Following acquisition of each MS2 spectrum, a synchronous precursor selection (SPS) MS3 scan was collected on the Top 10 most intense fragment ions in the MS2 spectrum. SPS-MS3 precursors were fragmented by higher energy collision-induced dissociation (HCD) at an NCE of 65% and analyzed using the Orbitrap.

Phosphoproteome analysis in heart tissues were processed with FAIMS/hrMS2 using our optimized workflow for multiplexed phosphorylation analysis on an Orbitrap Eclipse mass spectrometer. Briefly, the Thermo FAIMS Pro device was operated with default parameters (inner and outer electrode were set at 100°C, yielding a FWHM between 10 V to 15 V and dispersion voltage (DV) was set at -5000 V). Each fraction was analyzed twice by the mass spectrometer, once with a method incorporating two CVs (CV= -45 and -70V) and again with three CVs (CV= -40V, -60V and -80V) using a 2.5h method having a gradient of 6% to 30% B [60].

Mass spectrometric data for phosphopeptides fractions in kidney tissue were collected on an Orbitrap Lumos mass spectrometer. Mass spectrometric data were collected in CID mode and then processed with FAIMS/hrMS2 using our optimized workflow for multiplexed phosphorylation analysis with a method incorporating two CVs (CV= -45 and -70V). Detailed parameters for MS2 and MS3 are embedded in the RAW files.

### Mass spectrometry data analysis

Mass spectra data were processed using a Comet-based pipeline. Spectra were converted to mzXML using a modified version of ReAdW.exe. Database search included all entries from an indexed Ensembl database version 90 (downloaded:10/09/2017). This database was concatenated with one composed of all protein sequences in the reversed order. Searches were performed using a 50ppm precursor ion tolerance for total protein level analysis. The product ion tolerance was set to 1.000 Da for MS3-based analysis and 50ppm for MS2-based analysis, respectively. TMT tags on lysine residues, peptide N termini (+304.207 Da for heart and kidney tissues and +229.163 Da for liver tissue), and carbamidomethylation of cysteine residues (+57.021 Da) were set as static modifications, while oxidation of methionine residues (+15.995 Da) was set as a variable modification. In addition, for phosphopeptide analysis, phosphorylation (+79.966 Da) on serine, threonine, and tyrosine were included as variable modifications. Peptide-spectrum matches (PSMs) were adjusted to FDR < 0.01. PSM filtering was performed using a linear discriminant analysis (LDA), as described previously, while considering the following parameters: XCorr, ΔCn, missed cleavages, peptide length, charge state, and precursor mass accuracy. For TMT-based reporter ion quantitation, we extracted the summed signal-to-noise (S:N) ratio for each TMT channel and found the closest matching centroid to the expected mass of the TMT reporter ion. For protein-level comparisons, PSMs from all three tissues were identified, quantified, and collapsed to a peptide FDR < 0.01 and then collapsed further to a final protein-level FDR < 0.01, which resulted in a final peptide level FDR <0.001. Moreover, protein assembly was guided by principles of parsimony to produce the smallest set of proteins necessary to account for all observed peptides. PSMs with poor quality, MS3 spectra with TMT reporter summed signal-to-noise of less than 100, or no MS3 spectra were excluded from quantification. We provide an estimate for the probability of correct localization for each phosphorylation site using AScore algorithm [61]. 84%, 89% and 90% of the quantified phosphopetide used for analysis have an AScore greater than 13 in heart, liver, kidney tissues, respectively (P<0.05). All the information were uploaded to figshare (https://figshare.com/projects/Multiomics_analysis_identifies_drivers_of_protein_phosphorylation/137673, Under the folder titled “Raw summary of protein and phosphopeptides quantitation” - siteQuant5100.csv). The mass spectrometry proteomics data have been deposited to the ProteomeXchange Consortium via the PRIDE partner repository with dataset identifiers PXD032843.

### Sample preparation for transcriptomics analysis

Livers, hearts, and kidneys were dissected from each CC mouse, flash frozen, and stored at - 80°C. Once all samples were collected, frozen tissues were pulverized in liquid nitrogen, divided into aliquots, and then sent to the Genome Technologies service (Jackson Laboratory) for RNA extraction and RNA-seq analysis.

Total RNA was extracted and purified using the MagMAX mirVana Total RNA Isolation Kit (ThermoFisher) and the KingFisher Flex purification system (ThermoFisher). Briefly, pulverized tissue samples were lysed in TRIzol (ThermoFisher Scientific), chloroform was then added to the TRIzol homogenate, and the RNA-containing aqueous layer was removed for RNA isolation, following the manufacturer’s protocol. RNA concentration and quality were assessed using the Nanodrop 8000 spectrophotometer (Thermo Scientific) and Total RNA Nano assay (Agilent Technologies).

Libraries were constructed using the KAPA mRNA HyperPrep Kit (Roche) following manufacturer’s protocols. Briefly, poly-A mRNA was selected from total RNA using oligo-dT magnetic beads, followed by RNA fragmentation, first and second strand cDNA synthesis, ligation of Illumina-specific adapters containing unique dual index barcode sequences for each library, and PCR amplification. Library quality was assessed using the D5000 ScreenTape (Agilent Technologies) and concentration measured with the Qubit dsDNA HS Assay (ThermoFisher). Finally, pooled libraries were sequenced on the NovaSeq 6000 platform (Illumina) using the S1 Reagent Kit v1, yielding 20-40M (target 30M) 1 × 100bp single-end (SE) reads per sample.

## QUANTIFICATION AND STATISTICAL ANALYSIS

### Filtration of peptides that contain polymorphism

Peptides that contain polymorphisms, *i*.*e*., coding variants, bias protein abundance estimation in genetically diverse samples because peptides with differing sequences are not quantified simultaneously. A mouse with an alternative allele with respect to the B6 reference mouse genome will have reduced intensity or even non-detection for the reference peptide. This bias could then be propagated to estimates of protein abundance or phosphopeptide abundance, which can either obscure the signal of a true QTL or induce a false local QTL as a flag of the polymorphism. Therefore, we removed all polymorphic peptides based on the genome sequences of the founder strains. We filtered out peptides with known polymorphisms in the 8 founder strains of the CC. This is a more stringent filter than was applied in our previous study of the liver data [25] and thus there are minor differences in the protein quantification Before filtering, 153,856 unique peptides were quantified across 28 TMTs. 6841 (4.4%) unique peptides were filtered out due to not being identical in all 8 strains.

For each phosphopeptide quantified, the sequences of the corresponding protein were extracted from the founder strain genomes. To ensure that polymorphic peptides did not drive phQTL signal, we required phosphopeptides to have sequences of three amino acids adjacent to both sides that were present in all the founder strain genome sequences.

### Peptide normalization and protein abundance estimation

Peptides, including phosphopeptides, were standardized within TMT batch. The intensity for each peptide *j* from sample *i* was scaled by the ratio of the maximum cumulative peptide intensity in the batch to the cumulative peptide intensity for individual 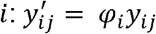 where

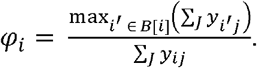

The abundance for each protein *j* from an individual *i* was estimated by summing the intensities of its component peptides (after removal of polymorphic peptides), then scaled relative to the abundance from the bridge sample and log transformed: 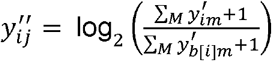 where *m* = 1, …, *M* indexes the peptides that map to protein *j* and 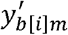 is the intensity of peptide *m* for the corresponding bridge sample from individual *i*’s batch.

We pre-adjusted for the effect of TMT batch using a linear mixed effect model (LMM) to allow strain pairs that span two TMT batches to be summarized to the strain level for downstream QTL analysis. The following LMM was fit for each protein *j*:

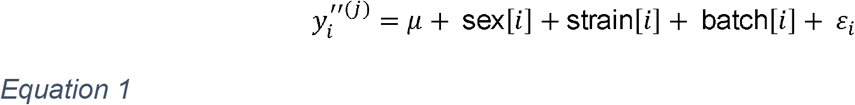

where 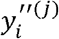 is the abundance of protein *j* for individual *i, μ* is the shared intercept, sex[*i*] is the contribution of sex for individual *i* (fit as a fixed effect), strain [*i*] is the contribution of the CC strain of individual *i* (fit as a random effect), batch [*i*] is the contribution of individual *i*’s TMT batch (fit as a random effect), and *ϵ*_*i*_ is the error for individual *i* with *ε*_*i*_ ∼ N(0, σ^2^) All downstream analyses were performed on quantities after subtracting off the batch effect: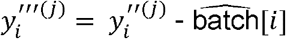, where 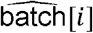 is the best linear unbiased prediction (BLUP) for the TMT batch of individual *i*. The LMM was fit using the lme4 R package^4^. Proteins that were unobserved for 50% or more of samples were removed from further analysis.

### Phosphopeptide normalization and adjustment for protein abundance

Phosphopeptides were processed similarly to protein abundance, but without the peptide-to-protein summation step. For each phosphopeptide *j*, we normalized as 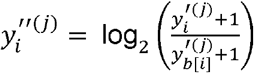 which were then batch adjusted as in Equation 1. Phosphopeptides that were unobserved for 50% or more of samples were removed from further analysis.

To distinguish genetic effects on phosphopeptides independent of the proteins from which they were derived, which we refer to as parent proteins, we also pre-adjusted for the effect of parent protein abundance by taking residuals from a linear model: 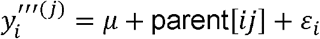, where parent [*ij*] is the contribution of the abundance of the parent protein for phosphopeptide *j* for individual *i* (fit as a fixed effect). The residuals for phosphopeptides were then calculated as: 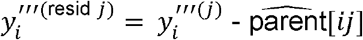. Parent protein abundances were estimated from their component peptides as previously described, but we first filtered out any peptides with a phosphorylation site.

### Heritability estimation

We estimated heritability for protein and phosphopeptide abundance in each of the tissues using an LMM. For a given protein or phosphopeptide *j* from a specified tissue, we fit:

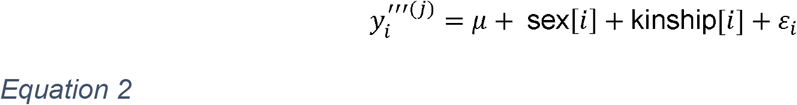

where kinship [*i*] is a random effect representing cumulative additive genetic effects and should thus capture similarities due to overall relatedness, modeled across individuals as **kinship** ∼ N(**0, G***τ*^2^). Bold text denotes vector and matrix quantities. **G** is a realized genomic relationship matrix, estimated from markers across all chromosomes, and *τ*^2^is the variance component underlying the kinship effect. Heritability is estimated as the proportion of variation due to genetic effects: 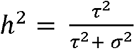. The qtl2 R package was used to fit the LMM and extract the heritability estimate^3^.

### Sex effects on protein and phosphopeptide abundance

Proteins and phosphopeptides that exhibited differential abundance between the sexes within a tissue, *i*.*e*., sex effects, were identified using the same LMM described in Equation 2 for heritability estimation. We instead compared it to a null model excluding the sex term and summarized the statistical significance with a likelihood ratio test *p*-value. The LMMs were fit using the qtl2 R package, using maximum likelihood estimates (MLE) rather than restricted maximum likelihood estimates (REML), as is appropriate for testing a fixed effect term. For a given outcome type (proteins or phosphopeptides), summary type (averages or differences), and tissue, significant sex effects were declared based on FDR < 0.1 using the BH method^5^.

### QTL analysis

For QTL analysis, we first summarized CC strain pairs as averages and differences (male – female) of the abundance of proteins and phosphopeptides. Phosphopeptides were adjusted for parent proteins as before, but at the strain level. We mapped QTL for proteins (pQTL), phosphopeptides (phQTL), and adjusted phosphopeptides (adj-phQTL). QTL for CC strain differences represent sex-by-genotype interactions where QTL effects differ between sexes. Founder haplotype probabilities for CC strain genomes were estimated by averaging the probabilities for the male and female at marker positions from MiniMUGA.

For each protein or phosphopeptide (unadjusted or adjusted) that was quantified in a tissue, we performed a genome-wide QTL scan by testing a QTL effect at positions across the genome.We fit a similar LMM to the heritability model in Equation 2 for each protein or phosphopeptide for a given tissue:

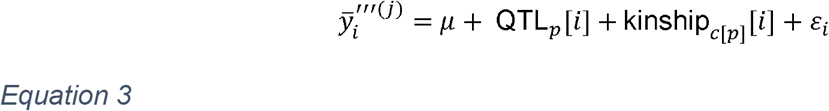

where 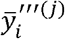 is the abundance summary (average or difference) for protein or phosphopeptide *j* of CC strain *i* and 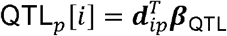 is the effect of putative QTL at marker *p* for CC strain *i* with 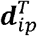 representing the founder haplotype probability vector for CC strain *i* at marker *p* (*e*.*g*., ordering the founder strains as AJ, B6, 129, NOD, NZO, CAST, PWK, and WSB, 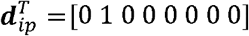 for a CC strain *i* that is B6/B6 at marker *p*), and ***β***_QTL_ is an eight-element vector of founder allele effects, fit as fixed effects. The kinship_*c* [*p*]_ [*i*] term is similar to the kinship effect in the heritability model (Equation 2), though instead modeled as kinship _*c* [*p*]_ ∼ N(**0, G**_*c* [*p*]_*τ*^2^), where the realized genetic relationship matrix **G** _*c*[*p*]_ used when testing markers as QTL on chromosome *c* is estimated by excluding all markers from chromosome *c, i*.*e*., the leave-one-chromosome-out (LOCO) method, to avoid the kinship term absorbing some of the QTL effect and reducing mapping power [62]. The kinship effect is used in mapping to account for potential population structure [63-66]. The strength of QTL significance was summarized by comparing the likelihood of Equation 3 to the likelihood of the null model excluding the QTL term, referred to as the log-odds (LOD) score. Mapping QTL for adjusted phosphopeptides is analogous to including the parent protein as a covariate in Equation 3 (and its null model). All genome scans for protein abundance and phosphopeptides were performed in the qtl2 R package [55].

### QTL significant thresholds

We estimated significance thresholds for QTL specific to individual proteins and phosphopeptides using permutations [67]. By performing 10,000 permutations for each outcome separately, outcome-specific thresholds account for differences in distribution, levels of missing data, and differing kinship effects. Permutation genome scans used the model in Equation 3, but with the data permuted by rearranging the CC strain labels on founder haplotype probabilities, thus breaking the QTL signal but not the kinship. We used a genome-wide error rate correction across marker loci and then applied an FDR correction to account for multiple testing across each outcome type within a tissue [68].

Specifically, for each outcome *j* within a tissue, we fit a generalized extreme value distribution (GEV) from the 10,000 maximum LOD scores from the permutations [69, 70]. For a given tissue and outcome type (strain averages or strain differences of proteins, phosphopeptides, and adjusted phosphopeptides), we calculated genome-wide permutation *p*-values as

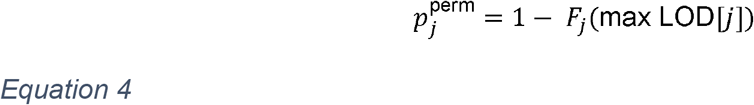

where *F*_*j*_ is cumulative density function for the GEV of outcome *j* and max LOD [*j*] is the maximum LOD score observed for outcome *j*. We then used the Benjamini-Hochberg (BH) procedure^5^ to calculate FDR *q*-values from the permutation *p*-values for a given tissue and outcome type. To estimate significance thresholds that are FDR-adjusted and outcome-specific, we applied interpolation to approximate permutation *p*-values to *q*-values < 0.1. Significance thresholds on the LOD score scale were then calculated as 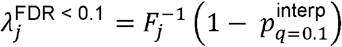, where 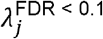 is the 10% FDR significance threshold for outcome 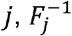 is the inverse cumulative density function for the GEV of outcome *j*, and 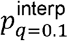 is the interpolated permutation *p*-value for *q*-value = 0.1.

### Defining local/distant status of QTL

As previously^1^, we defined detected QTL as “local” if their genomic coordinates were within 10 Mbp upstream or downstream of the middle of the coding gene and “distant” otherwise. We use the local/distant terminology instead of *cis*/*trans* because our definition is defined entirely by position and not genetic mechanism (*e*.*g*., *cis* regulatory elements). We used the broad 10 Mbp local window is broad, but the CC genomes have larger LD blocks than highly recombinant populations, such as the related Diversity Outbred population. Furthermore, we compared the effects of aligned QTL across tissues and sought to avoid aligned QTL being defined as local in one tissue but distant in another. However, the broad local window could misclassify some *trans*-acting QTL as local.

### Consistency of QTL across tissues

We evaluated the consistency of matched QTL (based on related outcomes and co-mapping to the same genomic region) by comparing their allele effects. We compared local and distant QTL across tissues (matching based on protein or phosphopeptide ID). We also compared co-mapping phQTL for unadjusted phosphopeptides to the corresponding local pQTL of the phosphorylation site’s parent protein for a given tissue. For matched local QTL, we only required them to be detected to be defined as co-mapping; for matched distant QTL, we also required that they were within 10 Mbp of each other.

Founder allele effects were estimated at the detected QTL marker, representing the ***β***_QTL_ term from the model in Equation 3. To stabilize the effects, they were modeled as a random effect: 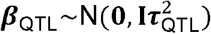, where **I** is the 8×8 identity matrix and 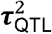 is a variance component underlying the allele effects. Allele effects were then estimated as BLUPs 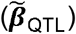, using the qtl2 R package^3^. The consistency of allele effects was summarized as the Pearson correlation coefficient between matched QTL: 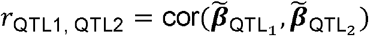, where QTL_1_ and QTL_2_ represent a co-mapping matched pair.

### Mediation of phQTL through parent protein abundance

We assessed whether detected phQTL were mediated through their parent proteins. For each phosphopeptide *j* with a detected phQTL in a specified tissue, we fit the following mode:

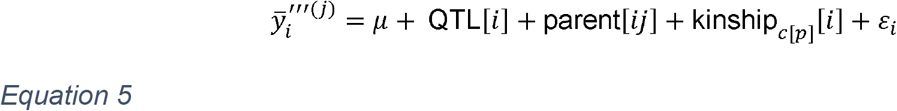

where 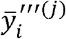 is the unadjusted abundance summary for phosphopeptide *j* with the phQTL for CC strain *i*, QTL [*i*] is as defined in Equation 3 but fixed at the peak marker for the detected phQTL being evaluated, and parent [*ij*] is the contribution of the abundance of parent protein of phosphopeptide *j* to strain *j*, modeled as a fixed effect.

We expanded the set of phQTL to include leniently detected ones (FDR < 0.5) for the evaluation of mediation through their parent proteins, providing a clearer picture of the large-scale mediation trends. A mediation LOD score for phosphopeptide *j* in a specified tissue, 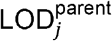, was calculated by comparing Equation 5 to a null model excluding the QTL term. To summarize across phQTL in a tissue, a Delta LOD was calculated by taking the difference between the mediation LOD score and the original LOD score of the phQTL. We note that a similar approach could be used to formally assess mediation of sex effects on phosphopeptides through parent proteins.

### Mediation of distant QTL

For each distant QTL detected in the CC tissues, we performed a mediation analysis analogous to the QTL genome scans [22, 24, 25, 71]. Instead of scanning through genetic markers as putative QTL, we scanned through putative mediators (from transcripts or proteins) of the specified distant QTL. A model similar to Equation 3 was fit:

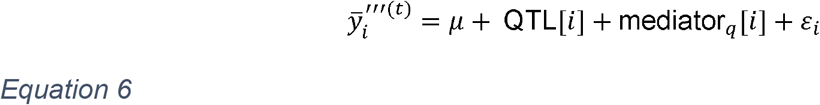

where 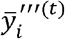 is the abundance summary (average or difference) for the target protein or phosphopeptide *t* with the distant QTL for CC strain *i*, QTL [*i*] is as defined in Equation 3 but fixed at the peak marker for the detected distant QTL, and mediator_*q*_ [*i*] is the contribution of the mediator *q* to individual *i*, fit as a fixed effect. The significance of the QTL term in Equation 6 is evaluated by comparing to the null model excluding the QTL term, producing a mediation LOD score: 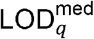. These summaries represent the distant QTL’s LOD score conditioned on each candidate mediator individually. We also note that the kinship effect is excluded from Equation 6 and its null model to simplify computation. Mediation scans were performed using the intermediate R package (https://github.com/churchill-lab/intermediate).

We assume that the vast majority of evaluated mediators for a specified distant QTL are not the true mediator, and thus the distribution of conditional LOD scores can be used as an empirical null distribution, approximately centered around the initially detected LOD score of the distant QTL. We calculate the *z*-scores of the conditional LOD scores and define strong candidate mediators as those with *z* < -8. Mediators are also expected to co-map a local QTL to the distant QTL. For distant pQTL, we evaluated proteins as mediators, whereas for distant phQTL, we evaluated both transcripts and proteins.

For candidate mediators highlighted in the Results, we estimated the strength of the relationships among QTL, mediator, and target based on proportion variance explained (PVE), calculated as 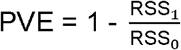. For the relationship between the QTL and mediator, RSS_1_ is the residual sum of squares from the QTL model (Equation 3) for the mediator and RSS_0_ is the residual sum of squares for the null model excluding the QTL term. For the relationship between mediator and target, the effect of the mediator on the target is evaluated rather than the QTL. We calculated a corresponding p-value for each relationship comparing the alternative and null models using ANOVA.

### Transcriptomics profiling

For each tissue, we aligned the RNA-seq reads using bowtie [72] to the pooled transcriptomes of the eight founder strains (Ensembl v84), and the alignments input to the genome reconstruction by RNA-Seq (GBRS) software to estimate total gene counts using EMASE. We used a variance stabilizing transformation [73] for the total gene counts for each tissue. As with protein and phosphopeptide abundance, the normalized expression for each gene was summarized at the CC strain level as averages and differences. Genes with no expression in 50% or more of samples were removed from further analysis. We also mapped eQTL using a similar approach as used for pQTL and phQTL, which we do not report here but make available at GSE199702.

## ADDITIONAL RESOURCES

All processed data and results are available for download and interactive analysis from the QTLViewer webtool (https://churchilllab.jax.org/qtlviewer/cc_phospho_peptides). Processed data and data analysis scripts have been deposited with FigShare (https://figshare.com/projects/Multi-omics_analysis_identifies_drivers_of_protein_phosphorylation/137673). The mass spectrometry proteomics and phosphoproteomics data have been deposited to the ProteomeXchange Consortium via the PRIDE [74] partner repository with the dataset identifier PXD032843. The raw transcriptome data have been deposited at GEO repository with the dataset identifier GSE199702.

## Acknowledgements

We thank the Genome Technologies service of the Jackson Laboratory for performing the RNA sequencing experiments. This work was supported by grant funding from the National Institutes of Health (NIH): K99CA273170 to T.Z.; F32GM134599 to G.R.K.; U19AI100625, P01AI132130 to F.P.-M.d.V. and M.T.F, R01ES029925 to F.P.-M.d.V.; R01GM067945 to S.P.G.; and R01GM070683 to G.A.C.

## Supplemental information

Supplemental tables

Table S1. Information about the 58 CC strains included in this study.

Table S2. Sex effect for proteins and phosphorylation sites in three tissues in CC strains.

Table S3. Heritability of proteins and phosphorylation sites in three tissues in CC strains.

Table S4. pQTL and phQTL summaries for three tissues in CC strains.

Table S5. Mediation analysis summaries for distant phQTL in three tissues in CC strains.

