## Supplemental Figures for "Multi-omics analysis identifies drivers of protein phosphorylation"

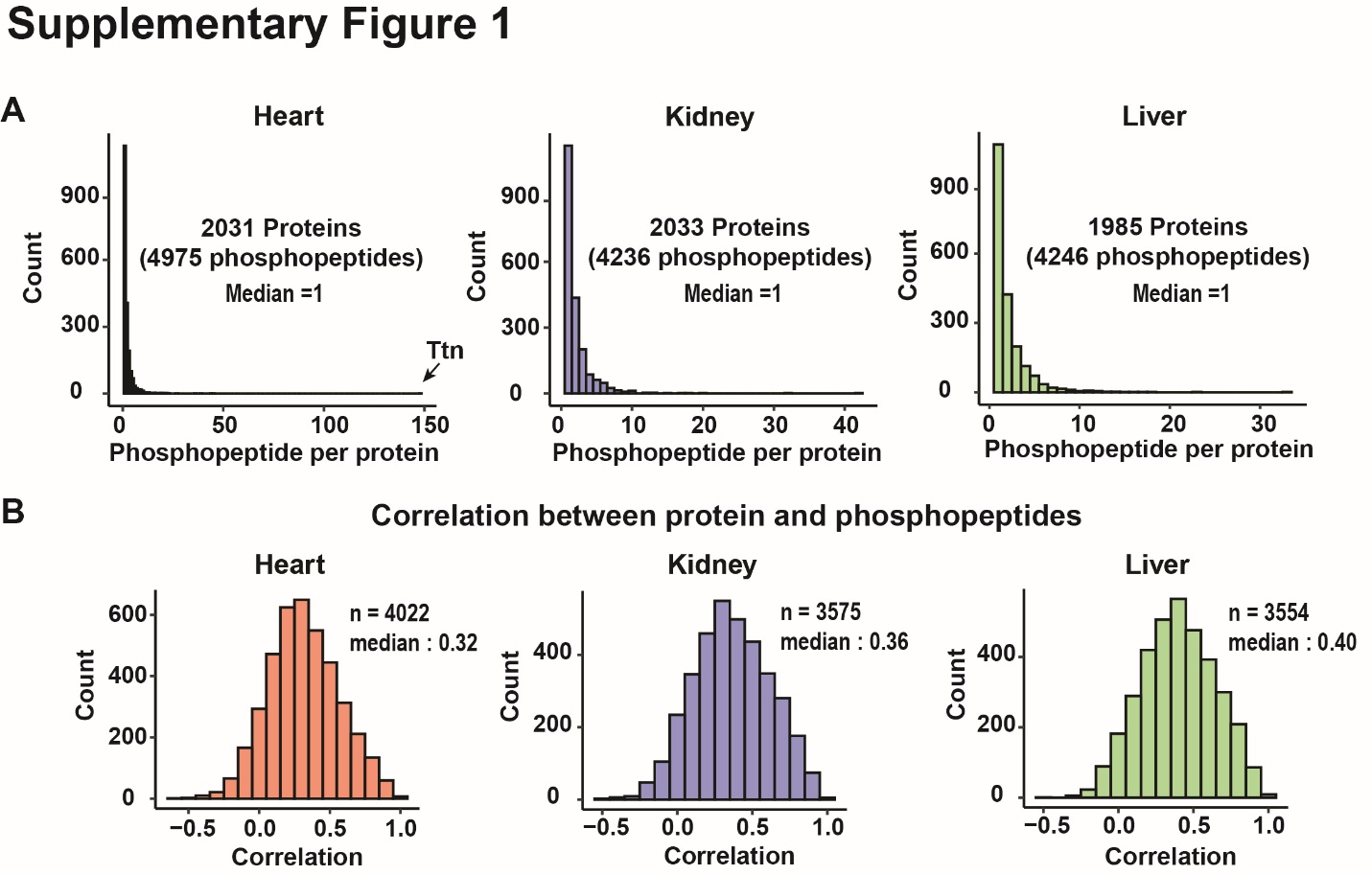

**Figure S1 (Related to Figure 1). Overview of protein and phosphopeptide quantification.** **(A)** Histogram of the number of quantified phosphorylation events per protein in three tissues. **(B)** Phosphopeptide abundances were highly correlated with their parent protein abundances in all three tissues. **(C)** Adjusted phosphopeptide abundances were not correlated with their parent protein abundances in all three tissues as expected.

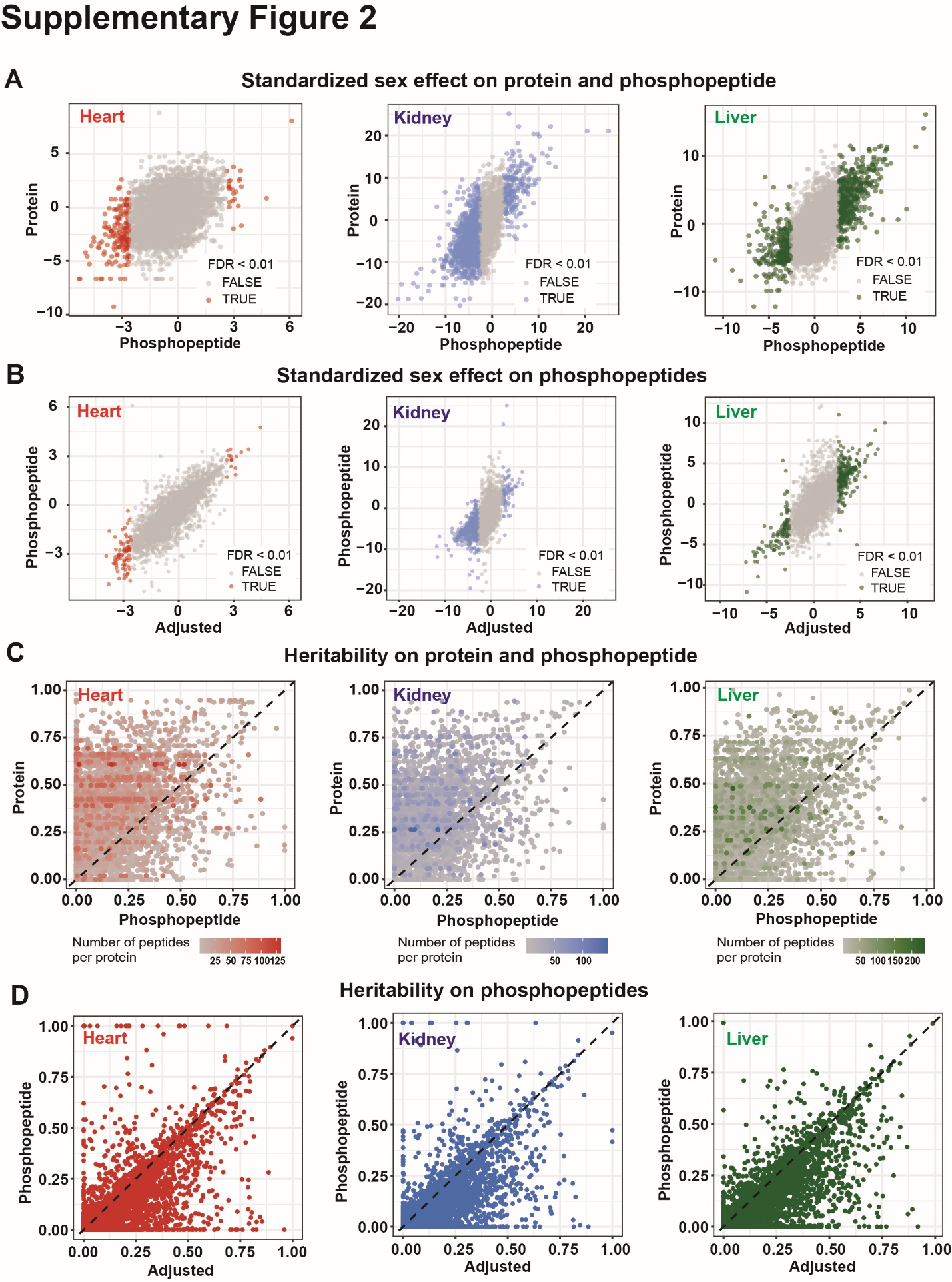

**Figure S2. Sex effects and heritability on the abundance of proteins and phosphopepitdes across tissues.** **(A)** Comparison of sex effects (difference/SE; female as reference) on phosphopeptide abundance and their parent protein abundance in heart, kidney and liver tissues. **(B)** Comparison of sex effect on phosphopeptide abundance before and after adjusting for parent protein abundance in heart, kidney and liver tissues. **(C)** Comparison of heritability on phosphopeptide abundance and their parent protein abundance in heart, kidney and liver tissues. **(D)** Comparison of heritability on phosphopeptide abundance and adjusted phosphopeptide abundance in heart, kidney and liver tissues.

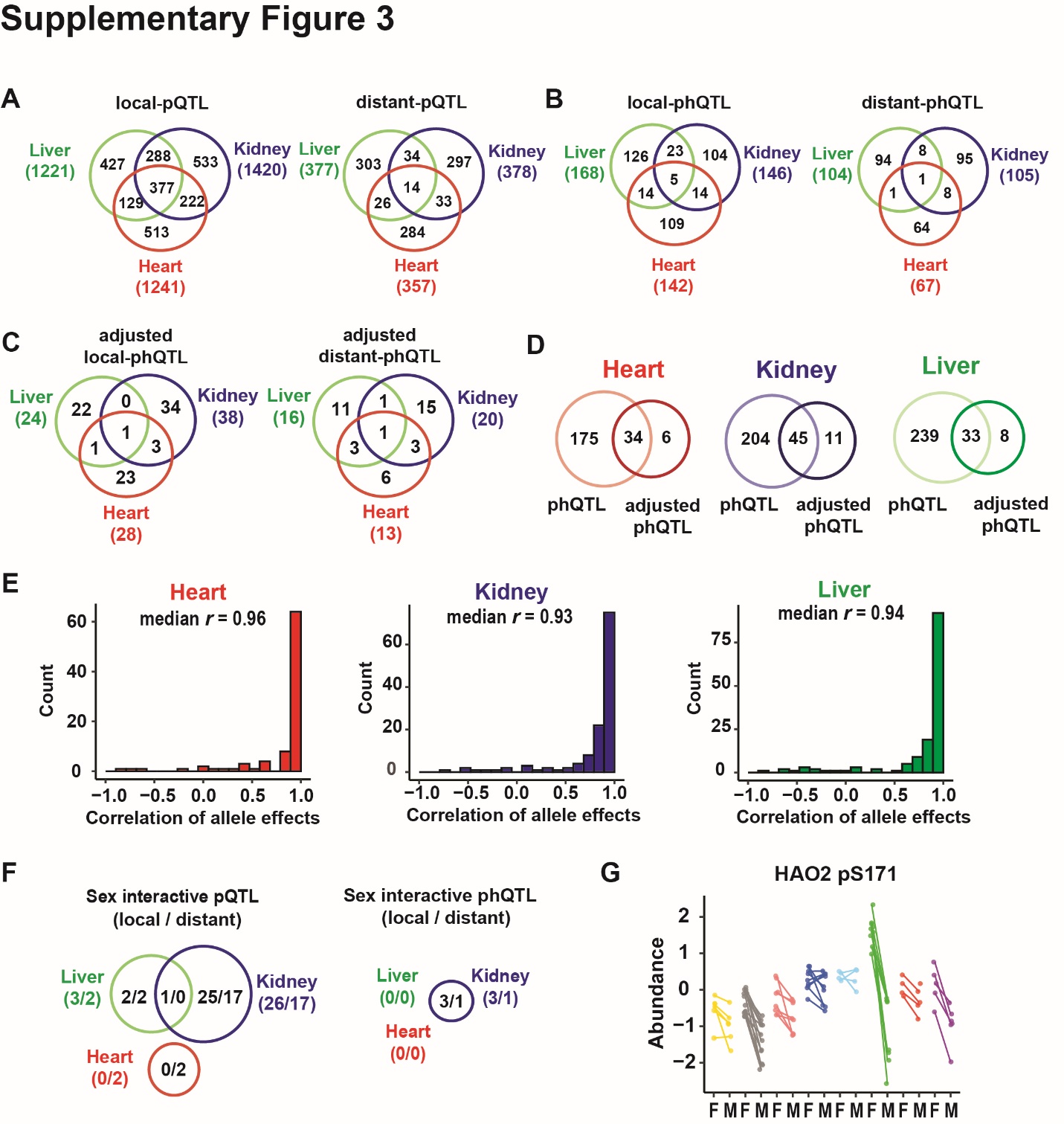

**Figure S3. pQTL and phQTL mapping from CC strains in heart, kidney and liver tissues**. **(A)** Venn diagram of the identified local pQTL and distant pQTL for proteins in liver, kidney and heart tissues. **(B)** Venn diagram of the identified phQTL for phosphopeptides in liver, kidney and heart tissue. **(C)** Venn diagram of the identified adjusted phQTL for phosphopeptides in liver, kidney and heart tissue. **(D)** Venn diagram of the identified phQTL and adjusted phQTL for phosphopeptides in liver, kidney and heart tissue, respectively. **(E)** The correlation of genetic effects for identified phQTL and pQTL in their parent proteins were high with exceptions in heart, kidney and liver tissues. **(F)** Venn diagram of detected sex-interactive pQTL and sex-interactive phQTL in three tissues. **(G)** HAO2 pS171 has a sex-interactive phQTL. Points are colored by founder haplotype at sex-interactive phQTL. Males and females from the same CC strain were connected by a line.

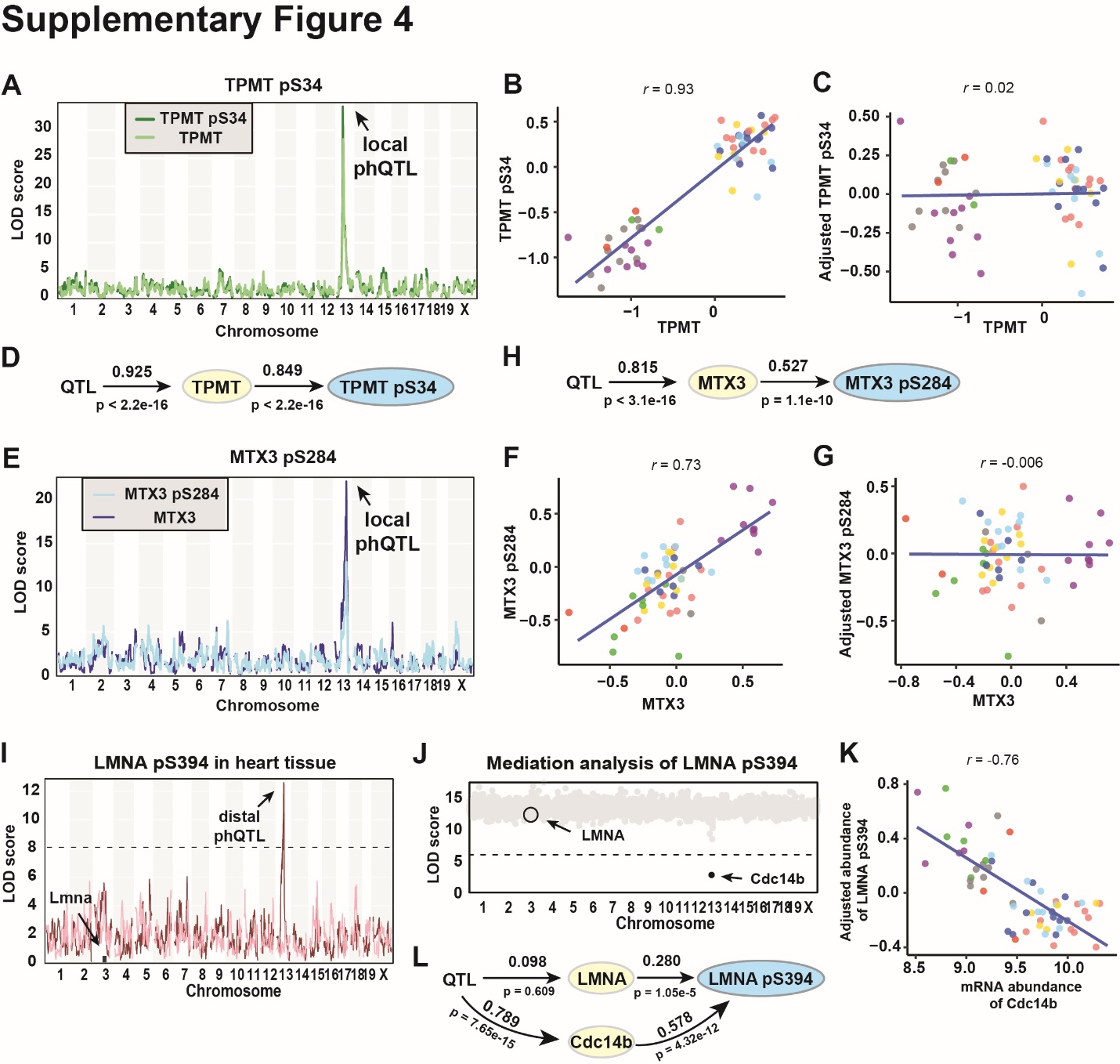

**Figure S4. Mediation of phQTL through the abundance of their parent proteins (substrates).** **(A)** Genome scans of TPMT and TPMT pS34 in liver tissue. **(B)** The abundances of TPMT and TPMT pS34 were highly correlated (*r* =0.93). **(C)** After adjusting for TPMT abundance, TPMT pS34 abundance is no longer correlated with TPMT abundance. Points are colored based on founder haplotype at *Tpmt*. **(D)** Path diagram of TPMT pS34 abundance regulation in liver tissue. **(E)** Genome scans of MTX3 and MTX3 pS284 in kidney tissue. **(F)** The abundances of MTX3 and MTX3 pS284 were highly correlated (*r* =0.73). **(G)** After adjusting for MTX3 abundance, MTX3 pS284 abundance is no longer correlated with MTX3 abundance. Points are colored based on founder haplotype at *Mtx3*. **(H)** Path diagram of MTX3 pS284 abundance regulation in liver tissue.

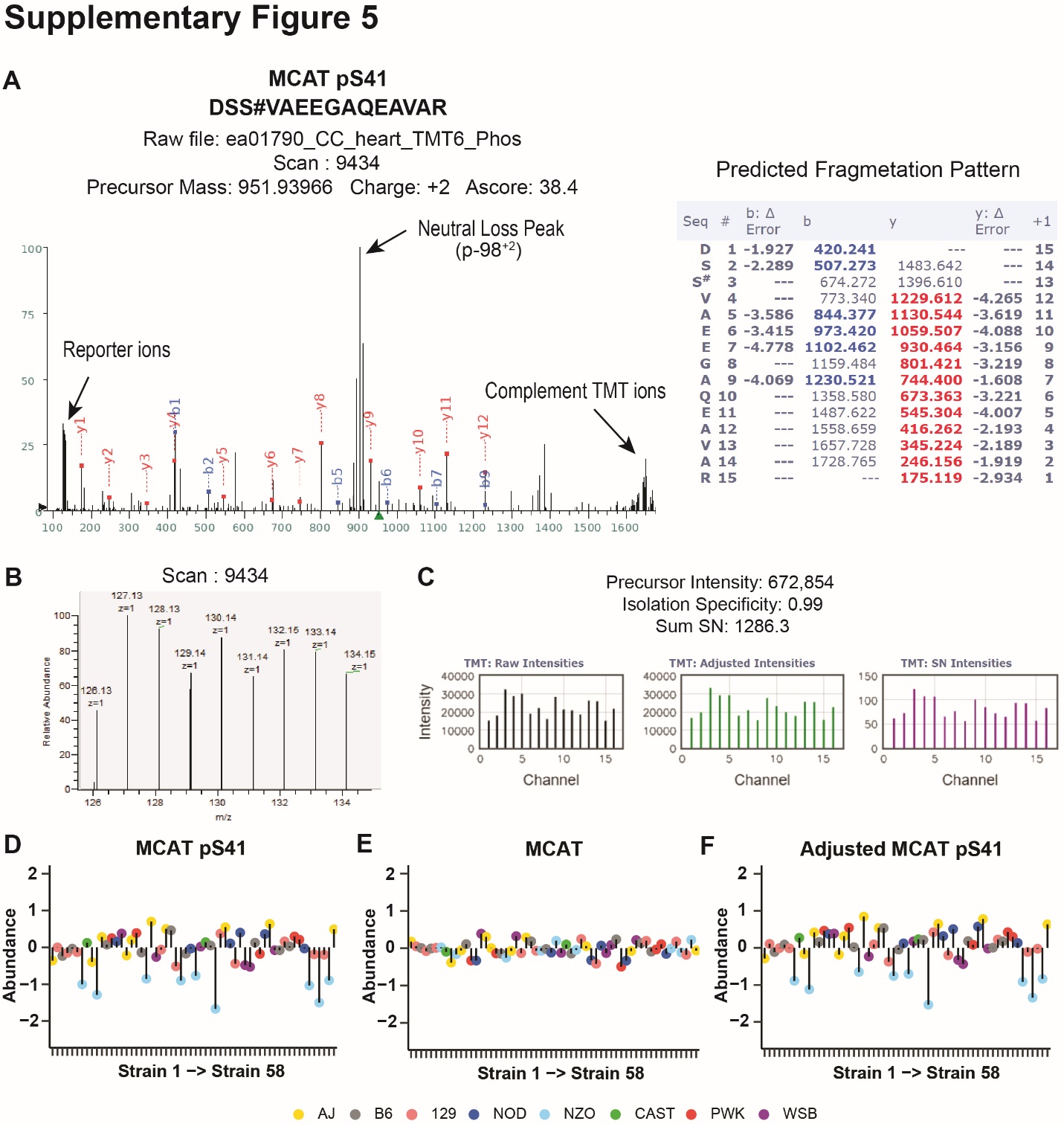

**Figure S5. *NZO* allele drives the low abundances of MCAT pS41 in CC strains.** **(A)** An example spectrum of identified phosphopeptide DSS#VAEEGAQEAVAR which harbors MCAT pS41. Match ions were highlighted in blue or red in the Predicted fragmentation table on the right. Reporter ions **(B)** of this spectrum was extracted and processed **(C)**. Signal to noise intensities were used for further analysis. **(D)**. *NZO* allele drives the low abundances of MCAT pS41 in 58 CC strains. Dots were colored based on the founder haplotye at the identified pQTL of MCAT pS41. **(E)** MCAT abundances have minimal variation in 58 CC strains. Dots were colored based on the founder haplotye at the identified pQTL of MCAT pS41. **(F)** *NZO* allele drives the low abundances of MCAT pS41 in 58 CC strains. Dots were colored based on the founder haplotye at the identified pQTL of MCAT pS41.

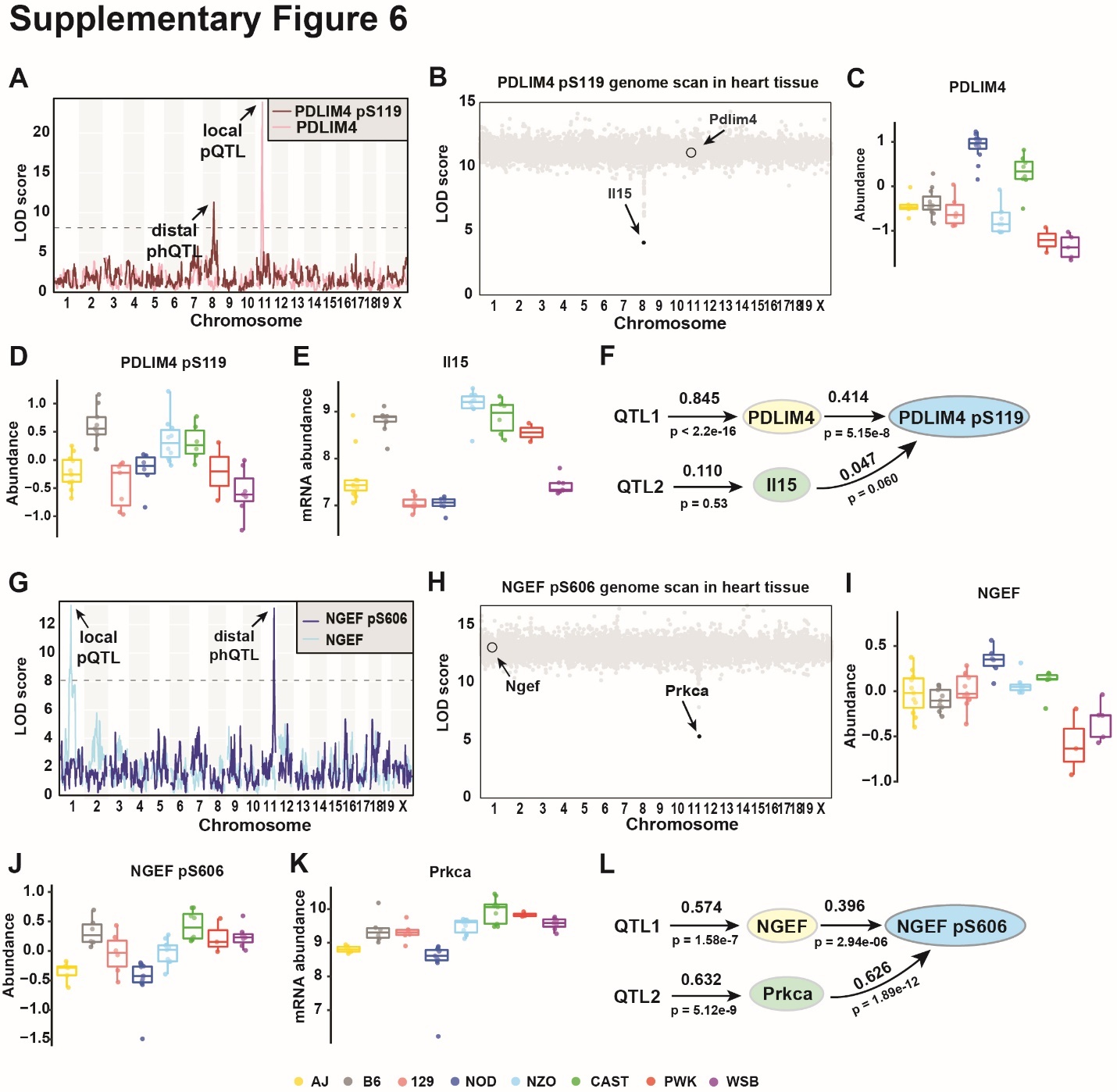

**Figure S6. pQTL and phQTL were identified to regulate phosphopeptide abundance together.** **(A)** Genome scans of PDLIM4 and PDLIM4 pS119 in heart tissue. **(B)** Mediation analysis using transcriptomics data identified *Il15* as the mediator for the phQTL of PDLIM4 pS119. Each gray dot is a mediation score representing the PDLIM4 pS119 LOD score conditioned on a transcript as candidate mediator. **(C)** The *NOD* and *CAST* alleles drove the high abundances of PDLIM4 in heart tissue. Data were categorized based on the founder haplotye at the identified pQTL. Abundances of (**D**) PDLIM4 pS119 and (**E**) *Il15* transcript had similar patterns based on the founder haplotype at *Il15*. **(F)** Path diagram of PDLIM4 pS119 abundance regulation in heart tissue. **(G)** Genome scans of NGEF and NGEF pS606 in kidney tissue. **(H)** Mediation analysis using transcriptomics data identified *Prkca* as the mediator for the phQTL of NGEF pS606. Each gray dot is a mediation score representing the NGEF pS606 LOD score conditioned on a transcript as candidate mediator. **(I)** The *PWK* and *WSB* alleles drove low abundance of NGEF in kidney tissue. Data were categorized based on the founder haplotye at the *Ngef*. Abundances of **(J)** NGEF pS606 and **(K)** *Prkca* transcripts had similar patterns. Data were categorized based on the founder haplotye at the *Prkca*. **(L)** Path diagram of NGEF pS606 abundance regulation in heart tissue.

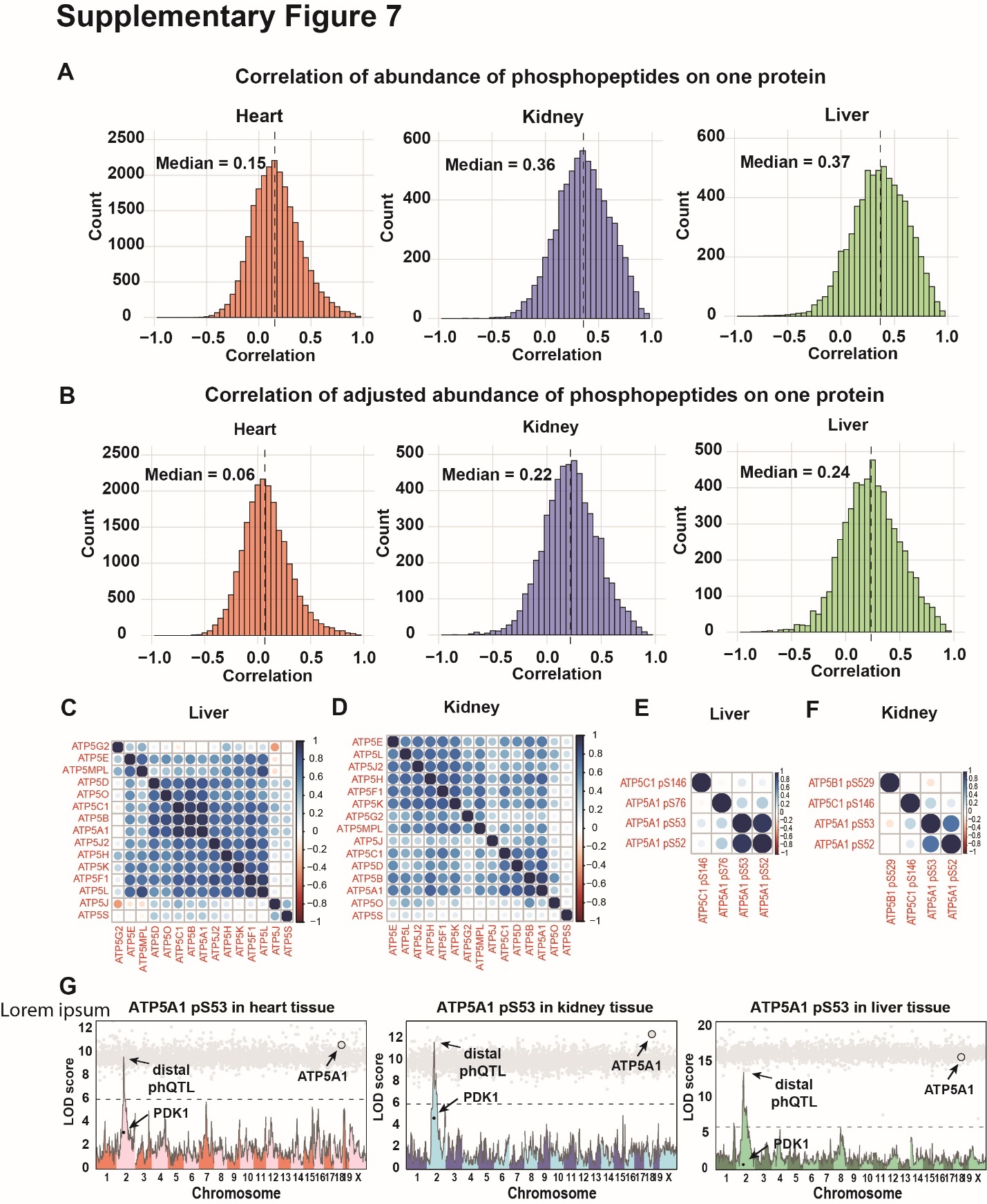

**Figure S7. (A)** Correlation of the abundances of phosphopeptides from the same protein in three tissues. **(B)** Correlation of the adjusted abundances of phosphopeptides from the same protein in three tissues. Protein abundance of subunits in ATP synthase complex were highly correlated in the **(C)** liver and **(D)** kidney tissues. Phosphopeptide abundance from the ATP synthase complex were not as correlated in **(E)** liver tissue and **(F)** kidney tissue. **(G)** Genome scans of ATP5A1 pS53 overlayed with mediation scores in (left) heart, (middle) kidney and (right) liver tissues. Each gray dot is a mediation score representing the ATP5A1 pS53 phQTL LOD score conditioned on a protein as candidate mediator.
